# Molecular orchestration of global actin remodelling by INF2

**DOI:** 10.64898/2026.08.14.744810

**Authors:** Felix Uecker, Sven Wargenau, Micaela Boiero Sanders, Lena Prange, Rafaella Jekabson, Annette Janning, Vanessa Krausel, Maximilian Gass, Hermann Pavenstädt, Daniela A. Braun, Michael P. Krahn, Christian Schuberth, Stefan Raunser, Peter Bieling, Roland Wedlich-Söldner

## Abstract

The actin cytoskeleton rapidly reorganizes in response to intracellular calcium signals, driving cellular morphogenesis and wound healing. Among actin regulators, the formin INF2 uniquely mediates the “Calcium-mediated Actin Reset” (CaAR) reaction, orchestrating transient and global actin remodeling upon calcium influx. Excessive INF2 activity is linked to kidney and neuronal diseases, underscoring the need for its tight control. Combining live cell imaging with single molecule tracking, biochemistry and structural analysis we discover that INF2 activity is tightly controlled by two interlinked mechanisms: canonical intramolecular autoinhibition and binding of the INF2 N-terminus to the side of actin filaments. Side-binding limits actin elongation and supports re-establishment of autoinhibition. Disruption of this negative feedback prolongs INF2 activity, affecting plasma membrane organization and repair as well as transcriptional control. Our findings uncover a novel product-inhibition mechanism that limits INF2 function and offer important insight into disease mechanisms linked to actin dysregulation.

## Introduction

The precise orchestration of actin dynamics drives fundamental physiological and pathological processes such as cellular morphogenesis, differentiation, wound healing and tumor metastasis^1,2^. Of particular importance is the capacity of the cellular actin cytoskeleton to rapidly reorganize and adapt in response to environmental cues or stresses^3^. Among the most ubiquitous and evolutionarily conserved cellular stress signals is a transient rise in intracellular calcium levels^4^. Calcium signalling controls most fundamental cellular processes, from cell energetics and secretion to muscle contraction and neurotransmission. Calcium signalling also acts directly on the actin cytoskeleton, regulating polymerization dynamics at multiple levels, including nucleation^5^, capping^6^, severing^7^, crosslinking^8^, and depolymerization^9^. Despite this central role, the molecular mechanisms that translate calcium influx into coordinated remodelling of entire actin networks are only beginning to be elucidated^10,11^.

A pivotal mediator of calcium-driven actin reorganization in mammalian cells is Inverted Formin 2 (INF2), a member of the formin protein family with the unique ability to respond directly to calcium signals^12,13^. INF2 is essential for the “Calcium-mediated Actin Reset” (CaAR), a highly conserved cellular program characterized by global and transient redirection of cortical actin into a cytoplasmic filament meshwork following calcium influx^14^. This stereotypical response occurs across diverse cell types and species. Misregulation of INF2 is linked to autosomal dominant forms of the kidney disease Focal segmental glomerulosclerosis (FSGS) and to Charcot Marie-Tooth neuropathy (CMT)^15^, underscoring INF2’s critical importance in health and disease.

Central to the CaAR mechanism is tight regulation of INF2. In resting cells, INF2 is kept inactive via intramolecular interaction between its Diaphanous Inhibitory Domain (DID) and Diaphanous Autoregulatory Domain (DAD), which flank the central FH1-FH2 domains responsible for actin filament nucleation and elongation^16^ (Fig. 1A). Although structurally similar to Rho-regulated Diaphanous-related formins^17^, INF2 is uniquely activated by calcium-bound calmodulin (CaM), which binds to a bipartite interface encompassing the N-terminal extension (NE) and the DID of INF2 to release autoinhibition^12,18^ (Fig. 1A), although this has not yet been studied in the context of CaAR. The mechanisms that spatially and temporally control INF2 activity during calcium signalling remain poorly defined. The prevalence of deregulated INF2 variants in disease contexts^15,19^ raise important questions about how cells ensure precise initiation - and more importantly - termination of INF2 activity.

**Figure 1.**
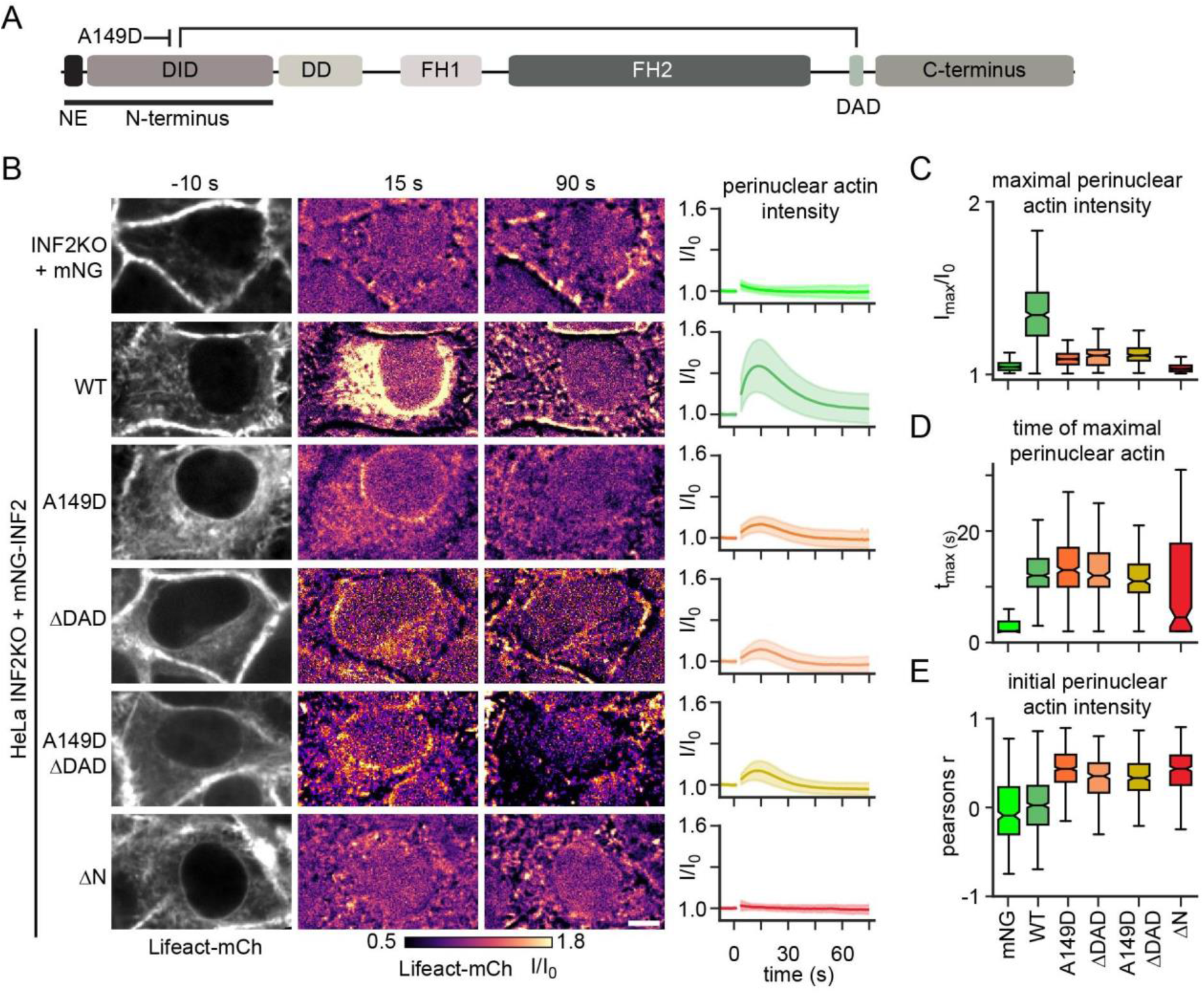
Regulation of INF2 activity by its N terminus. **(A)** Schematic representation of INF2 domains. NE: N-terminal Extension, DID: Diaphanous Inhibitory Domain, DD: Dimerization Domain, FH1/FH2, Formin Homology domain 1/2, DAD: Diaphanous Autoregulatory Domain. Intramolecular autoinhibition and its block by the A149D mutation is shown. (B) Actin reorganization by INF2 upon calcium stimulation. HeLa INF2 KO cells expressing Lifeact-mCherry and indicated mNG-INF2-CAAX constructs (or mNeonGreen - mNG as control) were imaged at indicated timepoints by spinning disk microscopy. Lifeact-mCherry images prior to uncaging of caged ATP are shown in grayscale. Images after stimulation show relative change in intensity (fire LUT) compared to the pre-stimulus condition. Graphs on the right indicate normalized Lifeact intensity profiles l/lo (Mean ± SD, n > 50, N = 3). (C, D) Boxplots for maximal levels of actin reorganization (Imax/IO, C) and the corresponding timepoint (tmax, D) for the cell lines shown in (B). **(E)** Box plots of linear correlation coefficients between Lifeact-mCherry and mNG-INF2 signals at t = 0 to describe basal/constitutive INF2 activity (n > 250, N > 3). Scale bar: 5 pm.

Here we combine quantitative live cell imaging with single molecule tracking in live cells, biochemistry and cryo-electron microscopy (cryo-EM) to uncover the mechanisms of INF2 regulation upon acute calcium stimulation and during CaAR. We identify multiple interactions within the INF2 N-terminus that orchestrate its activity cycle. We show that, beyond canonical autoinhibition, the N-terminal region of INF2 mediates actin filament side-binding, leading to product inhibition that limits filament elongation and facilitates formin re-inhibition. Defects in INF2 product inhibition impact plasma membrane (PM) wound repair, alter transcription via the Myocardin-Related Transcription Factor/Serum Response Factor (MRTF/SRF) pathway, correlate with disease-associated INF2 variants and perturb PM organization of Drosophila nephrocytes. Our findings provide novel molecular insights into INF2 regulation, describe for the first time the entire activity cycle of a formin in vivo and open new avenues for therapeutic intervention in conditions linked to aberrant actin homeostasis.

## Results

### INF2 regulation through calcium-calmodulin

To dissect INF2 regulation, we used the CaAR reaction as a quantitative readout^14^. INF2 knockout HeLa cells expressing Lifeact-mCherry (mCh) were transfected with mNeonGreen (mNG)-tagged INF2 variants of the ER-localized CAAX-isoform of INF2^20^, and calcium influx was triggered by uncaging of extracellular ATP (Fig. S1A-C). This induced the characteristic CaAR response, a rapid loss of cortical actin accompanied by a transient rise in perinuclear actin (Fig. S1G). We quantified the peak of relative increase in perinuclear actin (Imax/I0) as a robust measure of INF2 activity (Fig. S1D-I).

As expected, INF2 knockout cells showed no perinuclear actin response to calcium, while wild-type (wt) INF2 produced a peak accumulation within ∼15 seconds (Fig. 1B-D, movie S1). The autoinhibition-deficient mutant A149D^14,21^, which disrupts DID-DAD interaction, displayed elevated basal perinuclear actin levels (Fig. 1E) - yet, surprisingly, still responded partially to calcium stimulation (Fig. 1B-D). An identical partial response was seen upon deletion of the entire DAD domain (ΔDAD) or a combination of both mutations (A149D ΔDAD), ruling out residual DID-DAD contacts as an explanation. By contrast, deletion of the entire INF2 N-terminus (ΔN) produced a fully constitutively active formin that was completely unresponsive to calcium (Fig. 1B-E). Together, these results show that canonical DID-DAD autoinhibition alone does not account for the full regulation of INF2. The persistence of calcium-dependent control in A149D and ΔDAD mutants, and its complete loss only upon N-terminal deletion, points to an additional inhibitory mechanism residing within the INF2 N-terminus.

To explore the mechanistic basis for this regulation we next tested whether the observed activation kinetics of INF2 during CaAR was exclusively driven by calcium. A synthetic peptide covering the N-terminal extension (NE) of INF2 was shown to directly bind to calcium-calmodulin (CaM) with low affinity^18^ and a recent preprint characterized an additional binding site for CaM within the INF2 DID core^12^.

These dual interfaces were also predicted by structural modelling using AlphaFold 2 (Fig. 2A, S2). Key residues in INF2 are W11 in the NE that is predicted to bind to the C-lobe of CaM^18^ and I92 in the DID core that interacts with the CaM N-lobe^12^ (Fig. 2A).

**Figure 2.**
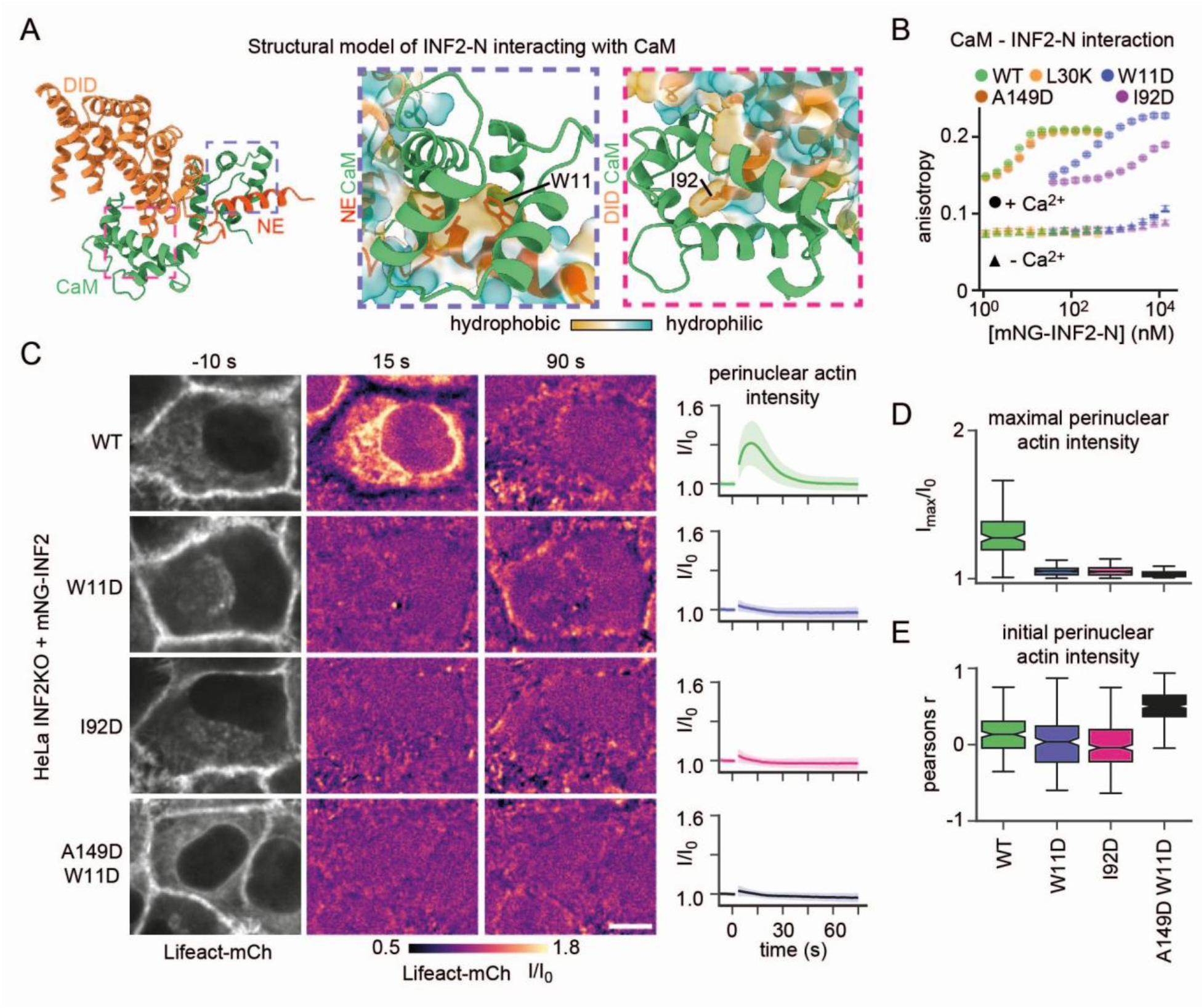
Activation of INF2 by CaM. **(A)** Alphafold 2 prediction of the interaction between the INF2N-terminus (INF2-N, orange) and calmodulin (CaM, green). Surface hydrophobicity and critical residues contacting CaM in INF2-NE (W11) and DID (192) regions are indicated in the two zoom-in boxes. **(B)** Binding of indicated recombinantly expressed mNG-INF2-N constructs and CaM-AlexaFluor594 in the presence and absence of calcium measured by fluorescence anisotropy (Mean + SD, N = 3). **(C)** Actin reorganization by INF2 upon calcium stimulation. HeLa INF2 KO cells expressing Lifeact-mCherry and indicated mNG-INF2-CAAX constructs were imaged at indicated timepoints by spinning disk microscopy. Lifeact-mCherry images prior to uncaging of caged ATP are shown in grayscale. Images after stimulation show relative change in intensity (fire LUT) compared to the pre-stimulus condition. Graphs on the right indicate normalized Lifeact-mCh intensity profiles in the perinuclear region I/lp (Mean + SD, n > 100, N > 3). **(D, E)** Boxplots for maximal levels of actin reorganization (D, Imax/lo) and pearson correlation coefficient of Lifeact-mCh and mNG-INF2 signals (E) at t = 0 for cell lines shown in (C). Scale bar: 5 pm.

To determine the contribution of either site to INF2-CaM complex formation we measured binding affinities between fluorescently labelled CaM and variants of the INF2 N-terminus via fluorescence anisotropy. WT INF2-N bound to CaM in a calcium-dependent manner with low nanomolar affinity (Fig. 2B, KD = 2.8 ± 0.7 nM). Consistent with the bipartite interface, the W11D mutant exhibited reduced affinity by two orders of magnitude (KD = 303.8 ± 10.7 nM) and the I92D mutant showed even lower affinity (KD = 4.28 ± 0.54 µM). Both CaM-binding mutants (W11D and I92D) effectively abolished activation of INF2 by calcium in cells (Fig. 2C, D). Importantly, the autoinhibition mutant (A149D) did not affect CaM-DID binding (Fig. 2B). Combining autoinhibition and CaM binding mutations (A149D + W11D) revealed the high basal activity expected from a lack of autoinhibition (Fig. 2E) but no additional activation by calcium influx (Fig. 2C-D).

In summary, we show that a high-affinity bipartite interaction between the INF2 N-terminus and CaM underlies the rapid activation of the formin by calcium. Importantly, the residual activation observed in the INF2 A149D mutant (Fig. 1B-C) is still dependent on CaM binding, indicating that calcium regulates the INF2 N-terminus in a manner distinct from the canonical DID-DAD autoinhibition.

### INF2 binds to the side of actin filaments via its N-terminus

To identify the molecular mechanism underlying this additional INF2 calcium regulation we expressed the isolated N-terminus (INF2-N) fused to mNeonGreen (mNG) in Hela cells. Strikingly, despite lacking the canonical formin F-actin binding domain (FH2), this fragment was clearly recruited to the cell cortex, where it colocalized with F-actin^15^ (Fig. 3A, S4G). Upon calcium influx, wt and A149D INF2-N rapidly lost cortical localization and redistributed to the cytosol (Fig. 3A). This response was blocked in the CaM binding deficient I92D mutant. These results suggest that INF2-N associates with cortical actin filaments in a calcium-sensitive, FH2-independent manner.

**Figure 3.**
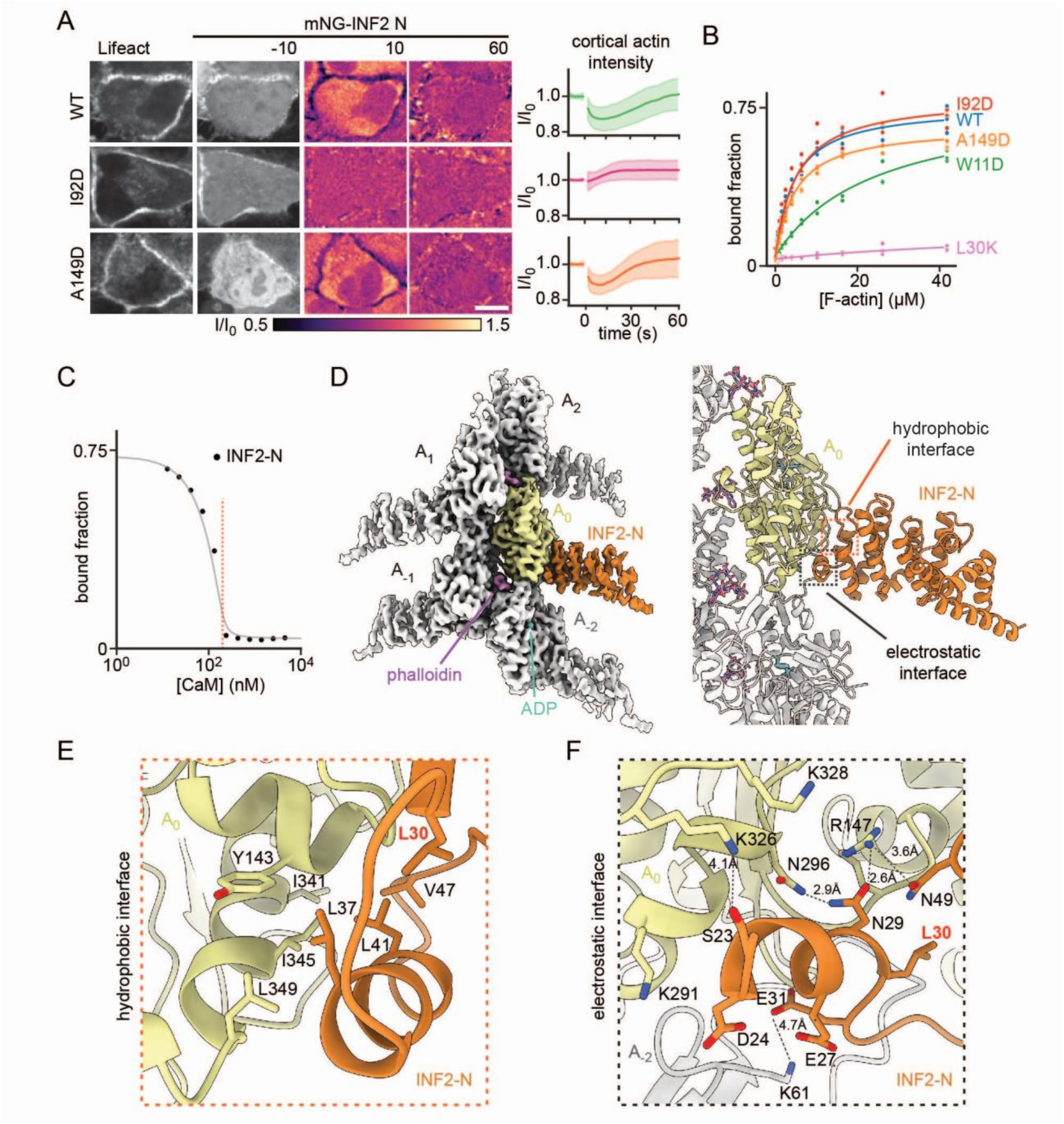
The INF2 N-terminus binds to actin filaments. **(A)** Exemplary images of Hela INF2 KO cells expressing Lifeact-mCh and indicated variants of the mNG-INF2 N. Graphs show the respective temporal cortical intensity profiles (n > 184, N = 3). **(B)** F-actin cosedimentation assays with bacterially expressed mNG-INF2-N variants (N = 3). **(C)** Inhibition of F-actin-lNF2-N co-sedimentation by CaM. **(D)** Cryo-EM density map at 3.48 A resolution (left) and molecular model (right) of F-actin decorated with INF2-N. **(E, F)** Zoom-in of hydrophobic **(E)** and electrostatic (F) interaction zones with key residues. Scale bar: 5 µm.

To directly test binding of INF2-N to F-actin we purified mNG-INF2-N variants and performed co-sedimentation assays with pre-assembled actin filaments. We found that both, wt and A149D INF2-N bound to F-actin with low micromolar affinity (Fig. 3B).

Structural information on the interaction of INF2-N with F-actin is missing. Therefore, to elucidate how exactly INF2-N interacts with actin filaments, we determined the cryo-EM structure of phalloidin stabilized INF2 DID-decorated β-actin filaments at a resolution of 3.48 Å (Fig. 3D, S3). The structure revealed that one INF2 DID specifically interacts with the interface between two consecutive actin subunits along the filament, resulting in a 1:1 molecular ratio in the F-actin-INF2-N complex. The general fold of INF2 DID resembles that of other formins that were determined in isolation by X-ray crystallography^22–25^. It comprises 4 armadillo repeats (ARM), and two post-ARM helices. In addition, it possesses a unique pre-ARM helix (α0) (Fig. S4A). The structure of the actin filament is similar to that of previously determined high-resolution F-actin structures^26,27^ (Fig. S4B), indicating that INF2 DID binding does not considerably alter the filament architecture. The DNase I-binding loop of actin (D-loop, residues 38-51), which is often not well defined in cryo-EM maps of F-actin, is highly ordered in our structure. It adopts an overall closed conformation, with the outer region displaced by 2.5 Å towards the filament centre compared to pure phalloidin stabilized β-actin filaments (Fig. S4B, movie S2). The closed D-loop position allows D34 of INF2-N to interact with G46 in the D-loop which would not be possible in an open D-loop conformation (Fig. S4B).

The high quality of the map at the binding site allowed us to model the side chains of all interacting amino acids. The region of INF2 DID involved in the interaction spans amino acids 23 to 49 and comprises two distinct binding interfaces (movie S3). The first interface is characterized by hydrophobic interactions between helix A1α1 (aa 35 to 42) and the cleft between actin subdomains 1 and 3 (SD1 and SD3) (Fig. 3E). This hydrophobic cleft interacts with many actin-binding proteins, including Lifeact and cofilin^28,29^ (Fig. S4C). The second interface spans residues 23-32 of INF2 (Fig. 3F). Several negatively charged amino acids of INF2-N form hydrogen bonds and polar interactions with positively charged amino acids in the vicinity of SD3 of the closest actin subunit (A0), and the D-loop of the actin subunit below it (A-2). Cofilin shows similar electrostatic interactions with this region^29^ (Fig. S4D). Our results indicate that INF2 dimers contains two N-terminal actin binding domains that individually bind to the side of filaments with moderate affinity. This binding could act as break for FH2-mediated elongation, while remodelling by cofilin is expected to displace INF2 from the side of filaments due to its high affinity^30^.

When comparing the actin-binding region of INF2-N with that of the CaM binding site we find significant overlap in their interfaces (Fig. S4E). A direct competition between the two binding partners for INF2 binding is consistent with the calcium-sensitive cortical association of INF2-N (Fig. 3A). Further validating a competition model, we found that in vitro interaction of the INF2-N terminus with F-actin was fully suppressed by stoichiometric amounts of CaM (Fig. 3C). Importantly, while the W11D mutation slightly reduced binding to F-actin, no change was observed for the I92D mutant (Fig. 3B). This suggests that F-actin and CaM compete for binding the INF2 N-terminus through partially overlapping binding interfaces.

However, we also identified residues that are uniquely involved in actin binding. In particular, INF2 L30 contributes to both actin-binding interfaces (Fig. 3E, F) but faces away from CaM (Fig. S4E). This makes mutations in L30 an ideal candidate for selectively uncoupling actin binding from CaM regulation. Indeed, an INF2 L30K substitution completely abolished F-actin co-sedimentation (Fig. 3B) as well as cortical localization (Fig. S4F, G) in cells, while leaving CaM binding fully intact (Fig. 2B; 4.1 ± 1.0 nM, n = 3). L30K thus serves as an ideal variant to characterize the contribution of actin side-binding to INF2 regulation.

To test whether actin side-binding contributes to INF2 inhibition in cells, we expressed the full length INF2 L30K mutant in INF2 KO cells and monitored the CaAR response. Strikingly, loss of side-binding produced an only slightly stronger, but markedly prolonged calcium-induced actin response (Fig. 4A, B), directly demonstrating that side-binding controls CaAR timing. A combination of the A149D autoinhibition and L30K actin side-binding point mutations led to constitutive cytosolic actin polymerization with no further response to calcium (Fig. 4A, B). This effect resembled a complete N-terminal deletion (Fig. 1B-E).

**Figure 4.**
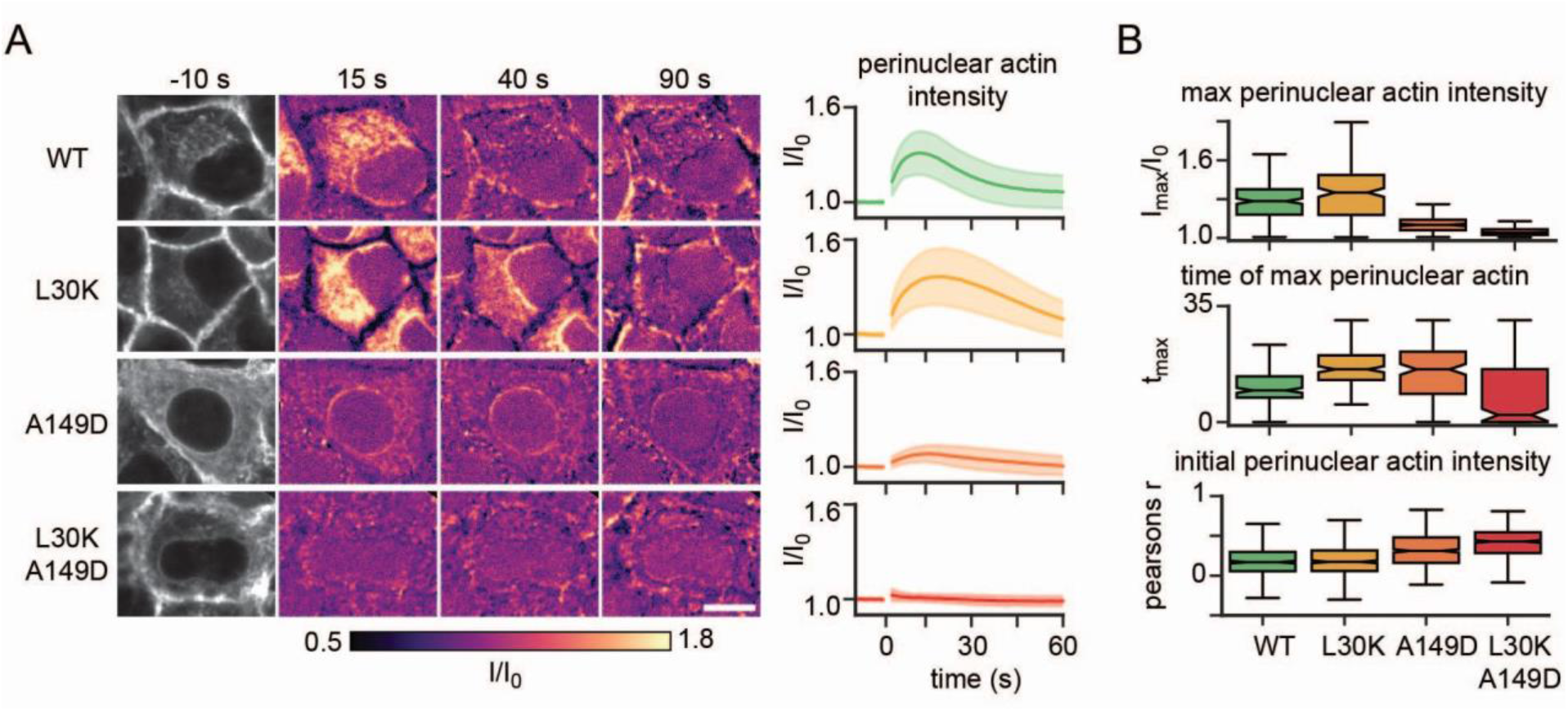
INF2 side binding regulates CaAR. **(A)** Actin reorganization upon calcium stimulation in HeLa INF2 KO cells expressing Lifeact-mCherry and indicated mNG-INF2-CAAX constructs (curves: n > 82, N = 3). **(B)** Boxplots for maximal levels of actin reorganization (lmax/lo). the corresponding timepoint (tmax) and correlation coefficient between Lifeact-mCherry and mNG-INF2 signals at t = 0 for the cells shown in (B) (n > 300, N = 3). Scale bar: 5 pm.

Other mutations that were predicted to selectively affect the actin side-binding interface produced the same effect (Fig. S4F-H).

Our results establish actin filament side-binding as a key regulatory mechanism for INF2 activity, independent of and in parallel to canonical DID-DAD autoinhibition. Actin side binding is key to rapidly shutting down INF2 activity following calcium stimulation.

### Actin side binding limits INF2 elongation and supports re-inhibition

The prolonged CaAR response of INF2 L30K could reflect two distinct, but not mutually exclusive mechanisms: side-binding might physically stall filament elongation, or it might facilitate re-establishment of DID-DAD autoinhibition. To distinguish these possibilities, we used single-molecule fluorescence microscopy to track individual INF2 dimers in living cells^31,32^, comparing variants deficient in CaM binding, elongation, side-binding, and autoinhibition (Fig. 5A).

**Figure 5.**
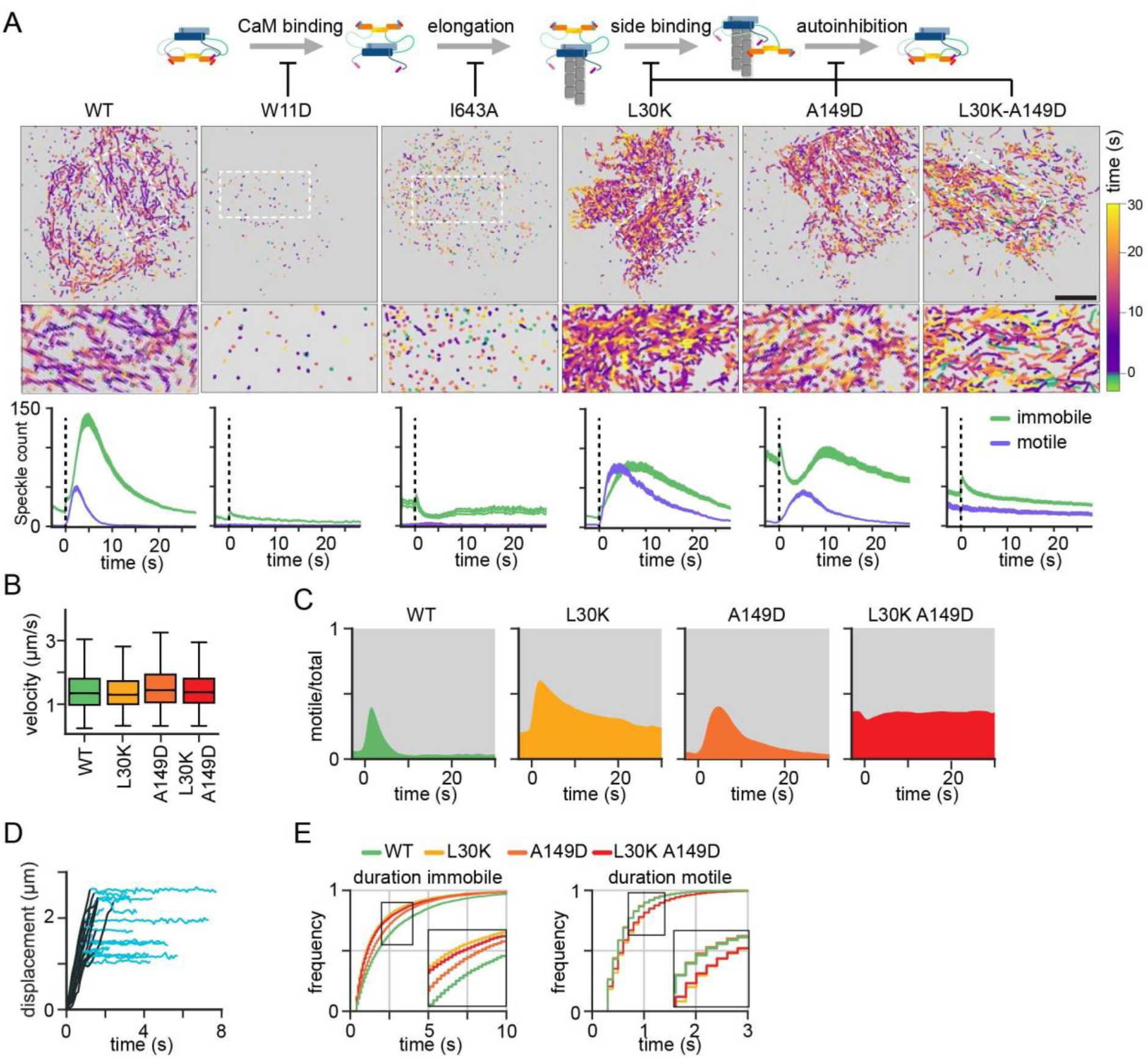
Single particle analysis of the INF2 activity cycle. **(A)** Tracking of single dimers of different NG-INF2-nCAAX variants. The schematic depicts the steps in the INF2 activity cycle affected by the respective variants. Detected particles are color-coded by the time of track appearance and projected (zoomed regions in white boxes). The graphs show mean and SD for total numbers of immobile and motile particles after calcium stimulation (t = 0, n > 150). Scale bar: 5 pm. **(B)** Box plots of velocities for motile particles of indicated INF2 variants (n > 10^5^). (C) Graphs showing INF2 activity after calcium stimulation for different regulatory variants. Activity is presented as the relative fraction of motile mNG-INF2 particles (n > 3500, N > 3). All variants were expressed at low levels in HeLa INF2 KO cells. (D) Plots of INF2 displacement for tracks from mNG-INF2 wt. Motile (black) and immobile (light blue) phases are indicated. (E) Cumulative frequency plots for the duration of immobile and motile phases of tracks for indicated INF2 variants (n > 5500).

Negative controls (CaM-binding deficient mutant W11D; elongation-deficient mutant I643A^33^) showed only a small baseline of immobile speckles and no motile speckles throughout (Fig. 5A). Upon calcium stimulation, both wt and L30K INF2 rapidly generated both motile and, with minor delay, immobile speckles (Fig. 5A, S5A, movie S3). Crucially, while wt motile speckles disappeared within ∼10 seconds, L30K motile speckles could be observed for more than 20 seconds after stimulation (Fig. 5A), matching the observed prolonged CaAR reaction in the absence of side-binding (Fig. 4A). Interestingly, the A149D autoinhibition mutant showed abundant immobile, but almost no motile speckles at baseline, which converted to motile speckles only upon calcium stimulation. This suggest that actin side-binding can safeguard INF2 activity even in the absence of canonical autoinhibition. The fully deregulated L30K-A149D double mutant on the other hand displayed a constant pool of motile speckles even without stimulation (Fig. 5A, S5A, movie S4).

Across all variants, processive elongation speeds during motile phases were indistinguishable (∼1.5 µm/s), indicating that the ability to bind to filament sides does not affect the speed of elongation once motility is initiated via Ca-CaM-mediated activation (Fig. 5B). Instead, normalizing the fraction of motile to total speckles revealed that the absence of side-binding in L30K and L30K-A149D dramatically extended the overall duration of INF2 activity (Fig. 5C).

The frequencies of motile and immobile particles in our single molecule experiments roughly corresponded to the actin remodeling kinetics observed in bulk CaAR assays (Fig. 4C). However, the prolonged activity of INF2 L30K was more evident at the single-molecule level, where motile INF2 L30K speckles persisted for more than four times longer than wt INF2 dimers (Fig. 5C). This difference likely reflects the limited cellular actin pool: actin polymerization may become saturated, so sustained INF2 activity does not result in proportionally greater actin assembly later during CaAR.

We next wanted to directly test a role for side binding in limiting INF2 elongation. Therefore, we used automated tracking routines to identify motile and immobile periods within INF2 tracks. To unambiguously assign INF2 activity states, we focussed on particles that initially appeared as motile, then halted before eventually disappearing from the plane of view (Fig. 5D). We found that the L30K and L30K-A149D variants exhibited longer motile phases than wt or A149D, indicating that actin side binding but not autoinhibition limits the processivity of INF2 (Fig. 5E). Interestingly, we find that duration of entire INF2 tracks (sum of motile and immobile phases) was limited to a few seconds irrespective of INF2 regulation (Fig. 5E). Therefore, physical stalling cannot explain the drastic difference in activity duration of wt vs L30K (Fig. 5C). Rather, INF2 L30K is able to polymerize multiple filaments before becoming non-motile and/or fully inhibited, showing a strong impact of actin side binding on the control of activity.

Taken together, our single-molecule data establish that actin side-binding constitutes the dominant mechanism controlling the duration of INF2 activity after calcium stimulation and can limit INF2 elongation, while DID-DAD mediated autoinhibition mostly determines steady state levels of INF2 activity.

### Differential control of INF2 activity via actin side binding and autoinhibition

Our single molecule tracking results suggest a crucial role of actin side binding, but not autoinhibition, in limiting the duration of INF2 activity. This predicts that mutations affecting INF2 autoinhibition should have only minor effects on the timing of its activity. Interestingly, a recent study showed that autoinhibition of the formin mDia1 proceeds through formation of a productive encounter complex between the DID and a charged stretch of the DAD domain^34^ (Fig. 6A). The corresponding sequence of the DAD in INF2 contains fewer charged residues (Fig. 6B), suggesting a weaker binding affinity. To test the consequence of this difference, we generated a series of INF2 variants with altered DAD charges (Fig. 6B). Consistent with a role for the charged stretch in mediating DID-DAD interaction, we found that increased positive charges in the DAD led to a strong reduction in CaAR amplitude, but had only marginal effects on CaAR duration (Fig. S6A, B). A corresponding charge decrease did not amplify the CaAR reaction, likely due to the saturation of overall actin levels in cells.

**Figure 6.**
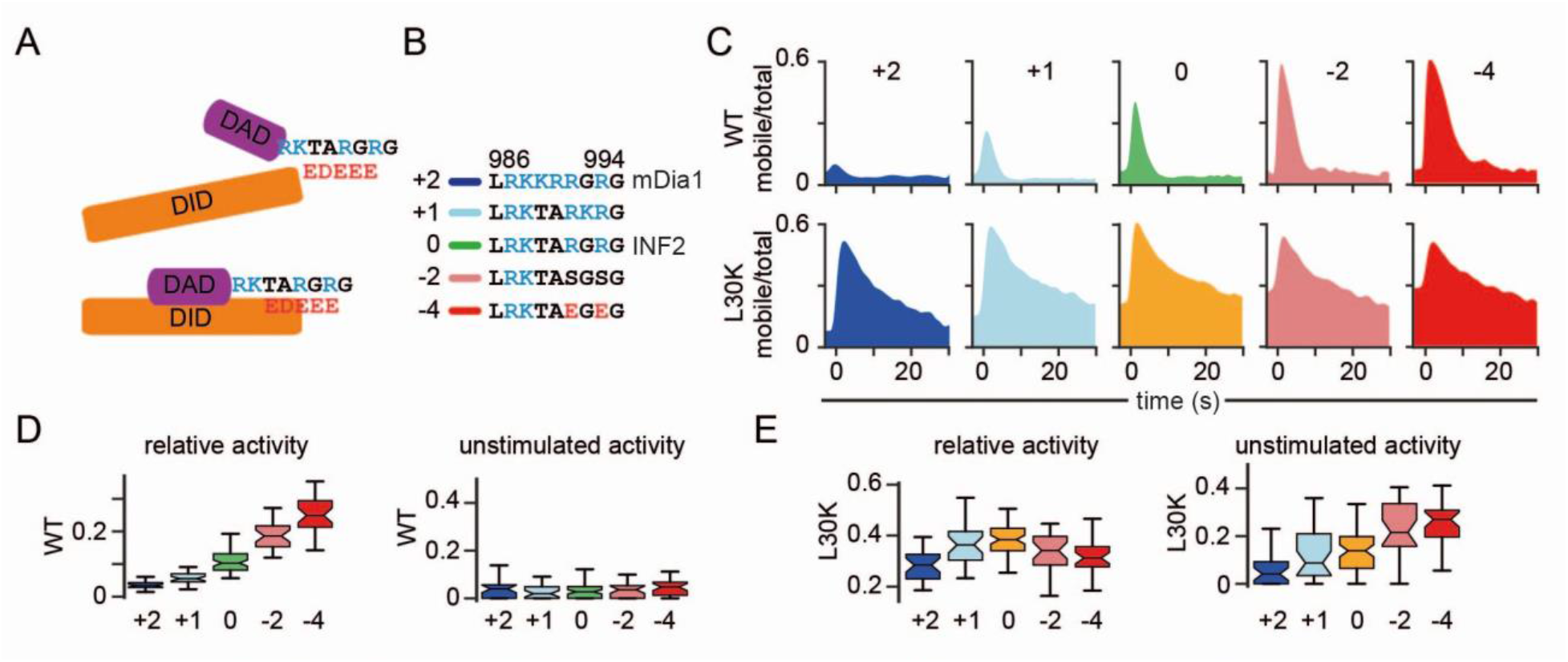
Actin side binding supports formation of an INF2 encounter complex. **(A)** Schematic representation of the charged region in the INF2-DAD that regulates formation of an encounter complex with INF2-DID. (B) Sequence alignement of the charged stretch and different INF2 variants used in this study. (C) Graphs showing INF2 activity after calcium stimulation for different DAD charge variants of INF2 wt and L30K, respectively. Activity is presented as the relative fraction of motile mNG-INF2 particles. (D, E) Box plots of relative activity (area under the curve in (C)) and unstimulated activity (activity at t = 0 in (C)) for indicated charge variants in INF2 wt (D) or L30K (E) (N > 29).

To assess whether DAD charge might nonetheless influence the duration of INF2 activity after calcium stimulation, we proceeded to examine the charge variants with our in vivo single molecule assay. We found a quantitative correlation between charge and the amplitude of relative INF2 activity (Fig. 6C, D, S6C). By contrast, the duration of relative INF2 activity was only moderately affected especially when compared with the actin-binding-deficient L30K variant (Fig. S6C). These findings indicate that DAD charge primarily regulates the probability or extent of INF2 activation, whereas it plays a lesser role in the timing of re-inhibition. In addition, basal activity remained low in all wild-type side-binding backgrounds (Fig. 6D), suggesting that steady-state inhibition was preserved. We next asked how the effects of actin side binding and DAD charge are integrated in INF2 re-inhibition. Notably, relative INF2 activity remained high in all L30K variants, regardless of overall DAD charge (Fig. 6C, E, S6D). In contrast, in the absence of side binding, unstimulated INF2 activity became strongly dependent on DAD charge (Fig. 6E). This was also reflected in the levels of F-actin in unstimulated cells (Fig. S6E,F).

Taken together, our findings demonstrate that actin side binding and DID-DAD interaction differentially regulate INF2 activity. Steady state levels of active INF2 in the absence of calcium stimuli are kept low via the DID-DAD interaction and the DAD charge primarily influences the probability of INF2-activation. In contrast, actin side binding is the main determinant of how rapidly INF2 returns to the autoinhibited state after calcium stimulation, thereby limiting the duration of activity. The two regulatory mechanisms combined ensure a rapid response to calcium, while maintaining low basal activity.

### Cellular role of dual safeguarding of INF2 via actin side-binding

Considering this dual safeguarding mechanism of INF2 activity, we wanted to determine how N-terminal regulatory interactions were linked to disease-relevant phenotypes. We previously reported that INF2 mutations linked to a combination of neurological and kidney pathologies (Charcot-Marie-Tooth disease: CMT and Focal segmental glomerulosclerosis: FSGS) induce formation of filopodia in HeLa cells^15^. In addition, corresponding variants of INF2-N were found to no longer localize to the cell cortex^15^. Indeed, we found that even without stimulation, expression of fully activated INF2 mutants, with side binding (L30K) and autoinhibition (A149D) abolished, led to strong formation of filopodia, with INF2 concentrated at their tips (Fig. 7A, S7A). In contrast, INF2-A149D, which still maintained actin side binding activity, did not induce formation of filopodia (Fig. 7A). To examine the cellular impact of INF2 upon acute activation we visualized cortical actin dynamics after plasma membrane wounding by laser ablation in INF2 wt vs. INF2 L30K expressing HeLa INF2KO cells^14^. We found that both variants supported formation of filopodia at wound sites, consistent with a temporal release of filament side binding for wt (Fig. 7B, movie S5). Importantly, while the number of filopodia was not significantly affected, filopodia formed by INF2 L30K were visible earlier (Fig. 7B) and became much longer (Fig. 7C). These results show that INF2 side binding activity not only changes the kinetics of actin polymerization, but also affects the architecture of the resulting cellular actin structures.

**Figure 7.**
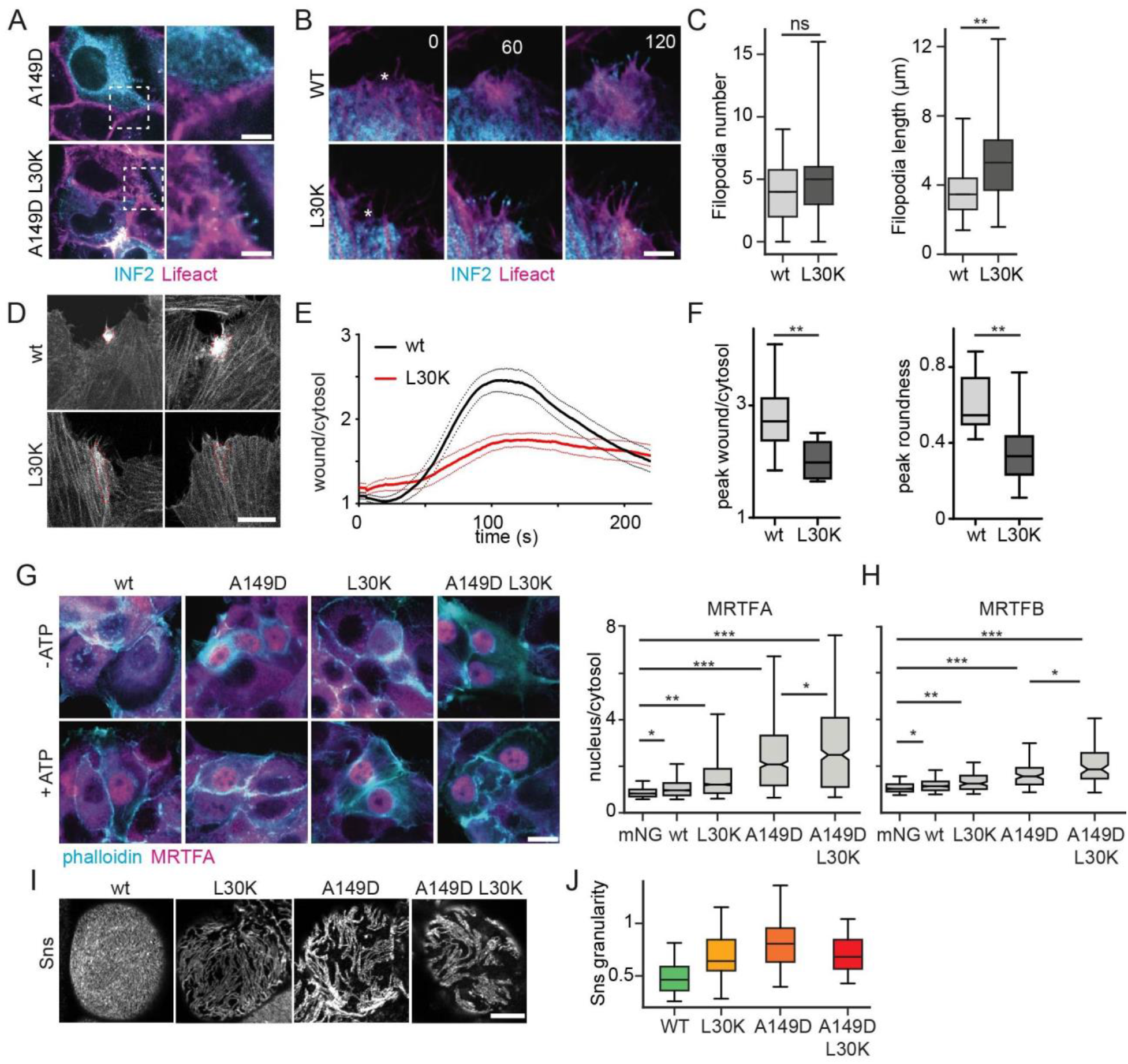
Physiological consequences of INF2 deregulation. (A, **B)** Formation of filopodia in HeLa INF2 KO cells expressing Lifeact-mCherry and indicated mNG-INF2 variants without stimulation (A) or at different times after PM wounding (B, wound positions indicated with asterisks). **(C)** Box plots of filopodia number and length for cells shown in (B) (n > 28). **(D, E)** Accumulation of actin at wound sites in HeLa INF2 KO cells expressing Lifeact mCherry and indicated mNG-INF2nC variants. Intensity profiles (E) show accumulation over time (n > 16, N > 3). **(F)** Box plots of amplitude and distribution for actin wound accumulation in (D) (n > 28, N > 4). **(G)** Immunofiuorescence images showing actin and MRTFA distribution in MCF7 INF2 KO cells expressing indicated mNG-INF2-CAAX variants without and with ATP addition. Box plots show corresponding nuclear MRTFA levels (n > 990, N > 3). **(H)** Box plots of nuclear MRTFB levels inHeLA INF2-KO cells expressing mNG-INF2-nCAAX variants (n = 370, N = 3) (I, J) Immunoflucrescence images of the fly nephrin Sns (1) and quantification of Sns granularity (J) in fly nephrocytes expressing indicated variants of Myc-INF2-CAAX (n > 68). Scale bars: 5 um.

The laser wounding assay enabled us to examine the consequence of INF2 side binding for large scale actin reorganization. A common feature of plasma membrane repair is a transient polarized accumulation of F-actin at the wound site^35^. We have previously shown that this accumulation depends on INF2^14^. Interestingly, while INF2 wt robustly supported wound accumulation of F-actin, cells expressing INF2 L30K exhibited a much weaker and less focused accumulation (Fig. 7D-F). These results suggest that rapid re-inhibition of INF2 via actin side-binding is required for efficient actin polarization towards the wound site.

We next tested whether the CaAR assay would allow us to quantify the contributions of side binding and autoinhibition to various disease variants. We therefore combined various FSGS and CMT linked variants with either the L30K or A149D mutation. One clear observation was that CMT variants are fully activated, similar to the A149D-L30K double mutant (Fig. S7B). This indicates that actin side binding is also affected in those disease variants. In contrast, all FSGS variants exhibited residual autoinhibition as they were able to respond to calcium stimulation and the combination with A149D consistently led to full activity (Fig. S7B). Interestingly, abolishing side binding in FSGS variants had variable effects, with E184Q-LK30K, S186P-L30K and L81P-L30K double mutants showing enhanced responses compared to the single disease mutants, no change for the Y193H and E220K and further reduction of CaAR response for L76P and L81P (Fig. S7B). In summary, our improved mechanistic understanding of INF2 regulation can help to further dissect the molecular basis for its associated diseases.

While in our single molecule experiments we found clearly increased unstimulated activity of INF2 L30K (Fig. 5C) this was not reflected in ensemble measurements of perinuclear actin (Fig. 4B), indicating that low constitutive activity of INF2 was not sufficient to affect overall actin levels in cells. We had previously shown that in MCF7 cells INF2-mediated actin polymerization during CaAR leads to the release of transcriptional co-activator MRTF-A from its inhibition by monomeric actin and its translocation into the nucleus, where it supports SRF-mediated transcription^14^. We reasoned that MRTF-A translocation could act as cumulative readout that was sensitive to low constitutive INF2 activity. Consistent with this premise, we found that expression of the CAAX-isoform of INF2 in MCF7 INF2KO cells led to a slight but significant increase in MRTF-A translocation in the absence of calcium stimulation (Fig. 7G).

In contrast, expression of the constitutive active INF2-A149D led to a strong increase of nuclear MRTF-A, which was further enhanced by the INF2 A149D L30K double mutant (Fig. 7G). Most importantly, expression of INF2 L30K led to a marked increase of MRTF-A translocation in the absence of stimulation (Fig. 7G), consistent with our single molecule measurements. We obtained similar results for translocation of MRTF-B when expressing the non-CAAX INF2 isoforms (Fig. 7H). These results indicate that even low constitutive activity of side-binding deficient INF2 can have consequences for gene regulation and cell physiology.

Finally, to validate the effects of INF2 deregulation in a pathophysiological context we overexpressed different INF2 variants in fly nephrocytes and observed its effects on distribution of the fly nephrin orthologue “Sticks and stones” (Sns). Consistent with our previous findings^15^, expression of wt INF2 did not affect Sns distribution, while constitutively active INF2 (A149D, A149D + L30K) induced a characteristic foot process effacements phenotype with reduced Sns density (Fig. 7I, J).

Importantly, expression of INF2 L30K was also associated with a significant Sns density reduction (Fig. 7I, J), comparable to our findings on weak FSGS-linked variants^15^. Our results indicate that loss of INF2 actin side binding can have far-reaching consequences on cell organization and can contribute to disease phenotypes linked to INF2.

## Discussion

Acute calcium elevation triggers a profound but highly transient reorganization of the actin cytoskeleton, a response linked to processes ranging from including plasma membrane repair, transcriptional control, and organelle homeostasis^14,36,37^. Although INF2 is a key driver of this response^13,14,38,39^, how its activity is turned on and off with such speed and precision has remained unclear. By combining quantitative live-cell microscopy, single-molecule imaging, biochemistry, and cryo-EM, we defined the full activity cycle of INF2 in vivo and discovered a set of two interlocking safeguards that ensure both rapid activation and timely shutdown (Fig. 8). Our data show that INF2 regulation cannot be explained by canonical intramolecular DID–DAD autoinhibition alone. Instead, calcium-bound calmodulin activates INF2 through a bipartite interface in the N-terminus, and this occurs in both wild-type INF2 and autoinhibition mutants. These findings agree with recent structural and biochemical studies describing the allosteric INF2 activation by CaM^12,18^. Importantly, we identify an actin filament binding site in the INF2 N-terminus site and show that this interaction is central for timely activity shutdown after calcium stimulation.

**Figure 8.**
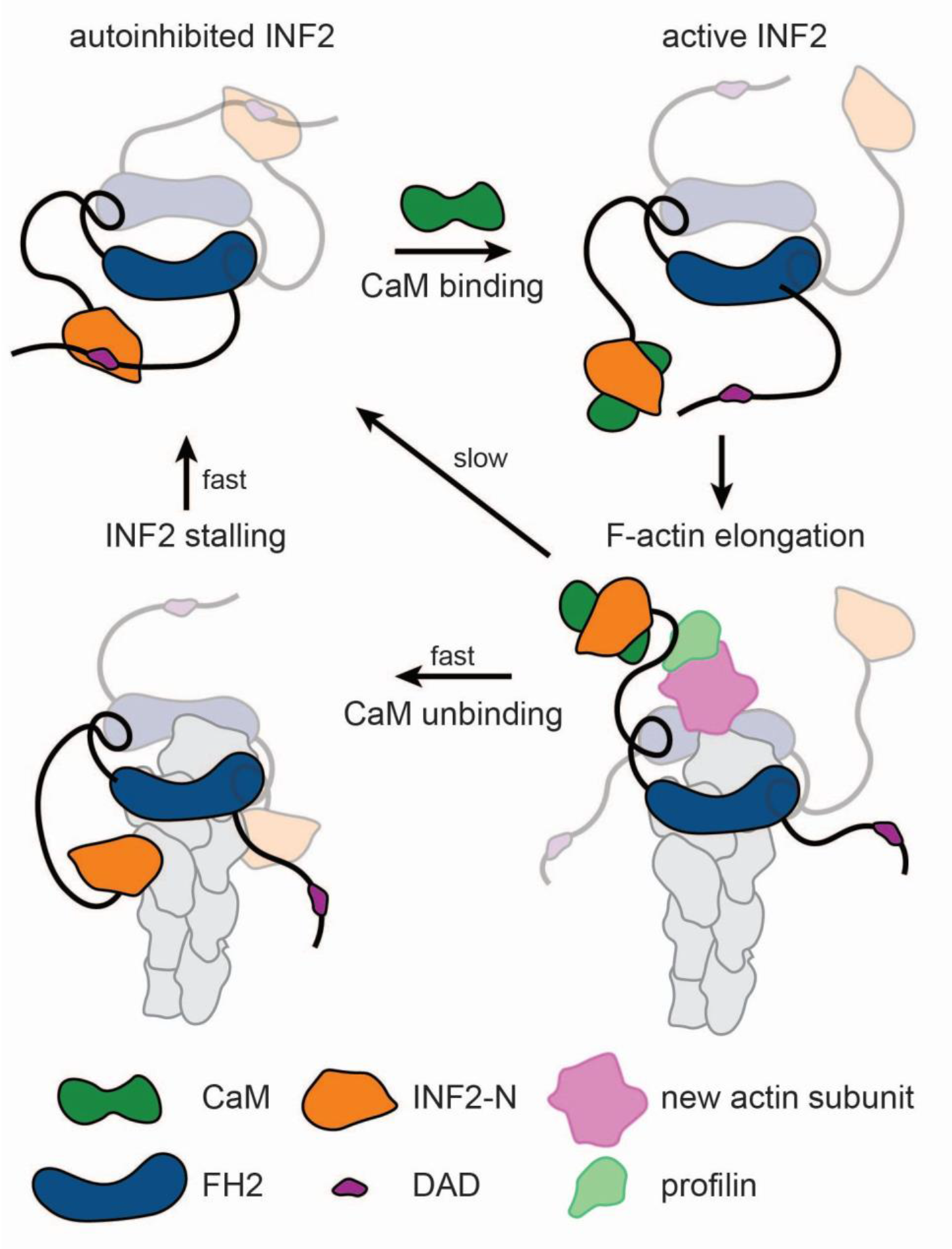
Schematic representation of the INF2 activity cycle The INF2 N-terminus binds to actin filaments. Binding of calcium-calmodulin (CaM) to the bi-partite interface in INF2-N disrupts INF2 autoinihibition by releasing the DID-DAD interaction. This allows the INF2 dimer to nucleate and elongate an actin filament via its FH2 domain. Once calcium is removed from the cell, CaM detaches and allows association of INF2-N with the side of the actin filament. This association stalls elongation and brings DID and DAD domains in close proximity, speedign up re-inhibition.

This side-binding mechanism is conceptually distinct from the monomer-sensing function of the DAD described previously^13^ and provides an intrinsic negative feedback that limits INF2-mediated elongation via product inhibition. Whereas G-actin binding promotes activation, F-actin side binding limits activity once polymerization has begun. INF2 therefore couples activation to product-dependent shutdown. Product-inhibition is a classical regulatory mechanism in many metabolic pathways^40^, but to our knowledge, INF2 represents the first example of an actin nucleator where binding to its own product filament provides negative feedback. The relevance of this mechanism is underscored by side-binding-deficient mutants, such as INF2 L30K and L30K-A149D, which exhibit substantially prolonged activity upon calcium stimulation (Fig. 5). These results indicate that side binding acts early during re-inhibition and is a major determinant of activity duration (Fig. 8).

Our findings support a model in which autoinhibition and actin side binding make distinct contributions to INF2 inhibition. DID-DAD interaction primarily sets the probability and overall level of activation, whereas side binding limits how long INF2 remains active after stimulation. This division of labor is supported by our DAD charge variants. Increasing DAD charge strongly reduced CaAR amplitude but had only modest effects on activity duration, whereas loss of side binding caused a pronounced prolongation of INF2 activity largely independent of DAD charge. Thus, autoinhibition mainly controls basal activity and activation strength, while actin side-binding provides rapid negative feedback needed for timely shutdown of activity.

The physiological consequences of this dual-safeguard regulation are substantial. In mammalian cells, even modest prolongation of INF2 activity disrupts spatial focusing of actin assembly and delays plasma membrane repair. Altered INF2 activity also changes MRTF/SRF-dependent transcriptional output, indicating that defective shutdown can affect gene expression well beyond the initial calcium pulse. Likewise, expression of side-binding-deficient INF2 in Drosophila nephrocytes causes defects in membrane organization and cell morphology, showing that these regulatory mechanisms are important not only biochemically but also in physiological settings.

More broadly, our work helps explain how cells achieve both responsiveness and restraint during calcium-triggered actin remodeling. INF2 must be activated rapidly and globally, but it must also shut down within seconds to prevent excessive polymerization. Product inhibition through actin side binding provides an elegant solution because it is intrinsically coupled to INF2’s own output. This makes the system robust to fluctuations in substrate availability and signaling noise.

Taken together, our study identifies INF2 as a formin whose activity is controlled by two complementary mechanisms: CaM-dependent activation and actin-side-binding-driven shutdown. This framework not only clarifies how INF2 supports fast and self-limiting CaAR responses, but also suggests more generally that regulated interactions with product filaments may be an underappreciated principle of cytoskeletal assembly. The strong sensitivity of cells to even small changes in INF2 activity duration further provides a mechanistic basis for understanding how subtle defects in INF2 regulation may contribute to disease. By defining the molecular basis for INF2’s activation and shutdown, our work provides a framework for targeted therapeutic strategies aimed at restoring actin homeostasis in human disease.

## Material and Methods

### Biochemistry

#### Actin purification for biochemical assays

Purification of β/γ-actin from bovine thymus was described previously^31^. Purified F-actin was stored at RT in F-buffer (2 mM Tris, 100 µM CaCl2, 200 µM ATP, 0.01% NaN3, 500 µM TCEP, 10 mM Imidazole, 50 mM KCl, 1mM EGTA, 1 mM MgCl2, pH7).

#### β-actin purification for cryo EM analysis

To obtain single isoform actin for structural analysis, human cytoplasmic β-actin was expressed as C-terminal fusion to thymosin β4 and a 10X-His-tag in insect cells^41,42^. A Cys-272Ala mutant was used to prevent oxidation in aqueous solutions. Purification was performed as described previously^43^. Purified G-β-actin was concentrated in a 10 kDa concentrator (Amicon) to 2 – 3 mg/ml, flash frozen in liquid nitrogen and stored at −80 °C.

#### INF2-N and calmodulin production

mNG-INF2-N variants (aa1-269) and Cys-Calmodulin1 (CaM) were expressed in BL21 Rosetta *E. coli* with TEV cleavable, N-terminal 6x His Tags (petMz2). Bacteria were grown in Terrific broth (Sigma Aldrich) at 37 °C to OD 0.8 before induction with 200 µM IPTG at 18 °C for 16 h. Cells were then resuspended in lysis buffer (75 mM NaPi, 350 mM KCl, 50 mM KAc, 0.4 mM β-mercaptoethanol,15 mM benzamidine, 1 mM PMSF, complete EDTA free protease inhibitors (sigma), 0.2 mg/mL DNAse I, pH 7.5) and lysed using a microfluidizer. Cell debris was removed via ultra centrifugation (200 000 g 45 min) and supernatants circulated three times over HiTrap™ columns loaded with 200 mM CoCl2. After thorough washing with wash buffer (75mM NaPi, 350 mM KCl, 50 mM KAc, 0.4 mM ß-mercaptoethanol,15 mM benzamidine, pH 7.5), proteins were eluted with a linear imidazole gradient (0-500 mM). Imidazole was removed via desalting (Cytivia HiPrep™ 26/10 Desalting column) and His tags were cleaved off with His-TEV-Protease on ice overnight. Finally, proteins were recirculated over CoCl2 columns to remove proteases and cleaned up via gel filtration (31.25 mM HEPES, 250 mM KCl, 62.5 mM KAc, 0.625 mM TCEP, pH 7.5). For storage, 20 % (w/v) glycerol was added and proteins were concentrated with Amicon concentrators. Samples were flash frozen in liquid N2 and stored at-80 °C.

#### Cryo-EM sample preparation of actin filaments decorated with INF2-N

Aliquots of recombinant human β-actin and mNG-INF2-N were cleared by spinning at 200,000 g in a TLA100 rotor for 30 min at 4 °C to remove aggregates. For mNG-INF2-N, storage buffer was exchanged to sample buffer (50 mM KCl, 1 mM EGTA, 1 mM MgCl2, 10 mM Imidazole, 10 mM KAc, 0.5 mM TCEP, 0.07% Tween). 40 µM β-actin was first incubated for 1 min in 10x ME (500 µM MgCl2, 2mM EGTA) to exchange ATP-associated divalent cation from Ca^2+^ to Mg^2+^ and then polymerized for 30 min at 37 °C by addition of 10x K50MEI (500 mM KCl, 10 mM EGTA, 10 mM MgCl2, 100 mM imidazole) and 40 µM phalloidin. Actin filaments were then pelleted at 100,000 g in a TLA100 rotor for 20 min at 4 °C, resuspended in sample buffer and mNG-INF2-N was added, with final concentrations of 8 µM actin and 60 µM mNG-INF2-N. The mix was used for cryo-EM preparation within 30 min.

#### F-actin co-sedimentation assay

Filamentous β/γ-actin was sedimented at 100 000 g for 30 min at 4 °C. Actin pellets were resuspended in Cosed-M buffer (1x KMEI, 5 mM β-mercapto-ethanol, 10 mM potassium acetate, pH 7). Actin concentration was determined at 290 nm. F-actin was stabilized with 1.2x molar excess phalloidin and 200 mg/mL Blocker™ Casein (Sigma Aldrich). Stabilized F-actin was serially diluted with Cosed-R buffer (1x KMEI, 5 mM β-mercaptoethanol, 10 mM KAc, 200 mg/L Blocker™ Casein, pH 7). Recombinant mNG-INF2-N variants were thawed and centrifuged in Cosed-M buffer at 200000 g for 30 min at 4 °C to remove aggregates. INF2 concentrations were adjusted to 600 nM in Cosed-R buffer. For binding analysis, 40 µL F-actin and 20 µL INF2-N were mixed for 10 min at 22°C. Samples were then centrifuged at 100 000 g, 4 °C for 30 min. Immediately after centrifugation, 40 µL supernatant was collected into black flat-bottom 96-well plates for measurement and remaining supernatant discarded. Pelleted protein was resuspended in 60 µL Cosed-R buffer, of which 40 µL were used for measurements. Fluorescence intensity was measured with a Tecan Spark plate reader at 22 °C with 470/10 nm excitation and 540/50 nm emission wavelengths.

The bound fraction was calculated with: 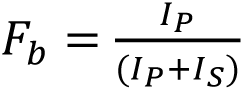

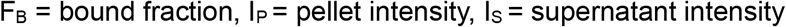

Resulting data triplicates were fitted globally^44^ with:

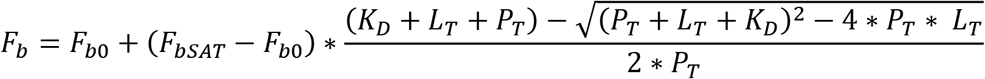

F_b0_ = bound fraction of background sedimentation, F_bSAT_ = bound fraction at saturating conditions, K_D_ = dissociation constant, L_T_ = total ligand concentration, P_T_ = total INF2-N concentration.

For competitive binding assays a fixed concentration and ratio of DID and filamentous actin was used and Cys-Calmodulin1 was added before the final centrifugation step.

#### Fluorescence Anisotropy measurements

To measure INF2-N - CaM interaction affinities, mNG-INF2-N variants were serially diluted and mixed with a constant concentration of 2.5 nM Alexa Fluor 594 maleimide-labelled CaM in anisotropy buffer (10 mM HEPES, 50 mM KCl, 1 mM MgCl2, 5 mM β-mercaptoethanol, 10 mM KAc, 200 mg/L Blocker™ Casein, pH 7) either with or without 0.35 mM CaCl2. Anisotropy measurements were performed with a Tecan Spark plate reader in black flat-bottom 96 well Corning plates at 22 °C with polarized excitation at 580/20 nm. KD values were obtained by fitting the following equation^44^:

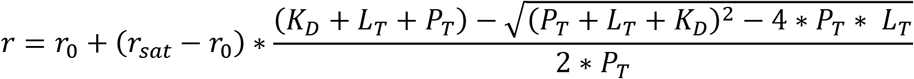

r_0_ = observed base anisotropy without INF2-N, r_sat_ = observed anisotropy at saturating conditions, K_D_ = dissociation constant, L_T_ = total INF2-N concentration, P_T_ = total AF594-CaM concentration.

### Cell culture and Molecular Biology

#### Plasmid construction

All mammalian INF2 expression plasmids used in this study were derived from a synthetic INF2 sequence (Twist Biosciences) based on the human INF2 isoform 1 (uniprot Q27J81) that included strategically placed restriction sites for easy domain swapping. INF2 was N-terminally fused to monomeric NeonGreen (mNG) separated by the following flexible linker sequence: “SGGGGSGGGGSGGGGSGGGGS”. To exchange between ER-bound (CAAX) and cytosolic (nC) INF2 isoforms, the plasmid was digested with KpnI and NheI followed by ligation with a 5’-phosphorylated oligo duplex with complementary 5’ and 3’ overhangs coding for the desired isoform. Similarly, DAD charge mutants were created by digestion with AgeI and BamHI followed by ligation of the desired mutant DAD domain encoded by a complementary oligo duplex. Site-directed mutagenesis of plasmids was carried out with KAPA HiFi Polymerase (Roche), according to the manufacturer instructions. Mutagenesis primers were created using the NEBasechanger tool. Template DNA was removed by 2 hour digestion with DpnI (NEB) prior to transformation into *E. coli*.

#### Cell culture and transfection

HeLa INF2KO cells stably expressing Lifeact-mCherry were described previously^14^. MCF7 INF2KO cells were generated via lentiviral transduction of INF2-specific gRNAs cloned into the lentiCRISPR v2 backbone^45^ and subsequent clonal selection on puromycin. Cells were cultivated in high glucose DMEM without L-glutamine supplemented with 2 mM L-glutamine (Capricorn), 1x non-essential amino acids (Capricorn), and 10 % fetal calf serum (Sigma-Aldrich), supplemented with 50 µg/ml hygromycin (InvivoGen). Cells were passaged by washing with PBS without calcium or magnesium and detached with 0.05 % trypsin/EDTA in DPBS (Capricorn) for 6 min at 37 °C. For transient transfection of cells, cells were detached and counted using the CellDrop 2 Channel Fluorescence Cell Counter (DeNovix). For transfection on the following day, cells were diluted to 8×10^4^ cells/ml or 6.5×10^4^ cells/ml for transfection after two days. 500 µl of cell suspension was seeded per well of an 8-well, glass-bottom slide (Sarstedt). For each well, we calculated the amount of plasmid DNA equivalent to 1×10E11 molecules of DNA using the theoretical molecular weight of the plasmid. For single molecule tracking experiments, we used 2.65 x10^10^ molecules. For transfection with more than 1 plasmid, equal numbers of each plasmids were transfected. Transfection was carried out using FuGENE6 transfection reagent (Promega) according to the manufacturer’s instructions.

#### Characterization of Drosophila Nephrocytes

Nephrocytes were analyzed as described before^15^. Fly strains were cultured on standard cornmeal agar food and maintained at 25°C. UASt::Myc-INF2 transgenes were established using the Phi-C31 Integrase system with attP40 as landing site. For overexpression of Myc-hINF2, virgin females of the nephrocyte-specific driver line sns::GAL4 were crossed with UASt::Myc-hINF2 males. Garland nephrocytes were isolated from wandering third-instar larvae by dissection, heat fixed, and stained using a chicken anti-Sns antibody (1:1000)^46^. Images were acquired on an SP8 confocal microscope (Leica).

#### Microscopy

For all live-cell imaging experiments, culture medium was replaced with 450 µl of HBSS containing calcium and magnesium (Biochrom), supplemented with 10 mM HEPES (Life Technologies). Imaging was performed at 20 °C on a custom iMIC microscope from Thermo/Till Photonics equipped with a 405/488/561/640 nm Sole-4 laser combiner (Lambert Instruments/Omicron) connected to spinning disk and Polytrope (epifluorescence and TIRF) beam paths via separate multimode fibers. Images were acquired on two iXon Ultra 897 EMCCD cameras (Andor) with no binning.

#### MRTF A/B immunofluorescence

A high-throughput automated imaging workflow was established to quantify MRTF nuclear translocation. One day before visualization of MRTF-A/B MCF7 cells were seeded onto 96 well glass bottom dishes and transfected with mNG-INF2 constructs. Cells were fixed with 4 % paraformaldehyde in PBS for 20 min, washed 3x with PBS and used directly for staining. Cells were permeabilized with 0.1 % TritonX-100 in PBS for 5 min and then washed 3x in PBS. Samples were blocked with 10 % goat serum for 30 min at RT. After washing 3x in PBS, primary antibody (anti-MRTF-A: mouse, Santa Cruz sc-390324, anti-MRTF-B: rabbit, Novus biologicals NBP1-46209) was added at 1:100 for 1h at RT, washed 3x in PBS. Fluorescent secondary antibody (goat anti-rabbit-Alexa Fluor 488, ThermoFisher A-11008 and goat anti-mouse Alexa Fluor 488, ThermoFisher A-11029) was then added at 1:400 for 1 h at RT. For F-actin staining Alexa Fluor 568 Phalloidin (Invitrogen A12380) was added to the secondary antibody mixture at 100 nM final concentration. Dishes were washed 3x in PBS and mounted with Moviol + Dabco.

#### cATP CaAR-assay

Calcium influx in cells was induced by photo-uncaging of nitrophenylethyl ester (NPE)-caged ATP (Thermo Fisher). Images were acquired with a 40x air objective (NA 0.9, Olympus). Photolysis of NPE was induced by scanning a 9 µm^2^ area with a 405 nm laser set to 100% intensity. The calibrated scanning point diameter was 0.22 nm, with 20% oversampling and a pixel dwell-time of 80 ms/µm^2^. While images were acquired in spinning-disk mode, photolysis required the epifluorescence beam path, which caused a 2 sec delay between the last pre-stimulation frame and the first post-stimulation frame due to filter changes. cATP was added to each well at a final concentration of 50 µM prior to imaging. At the start of each movie, a reference image was taken to capture the transfected mNG-INF2 construct, Lifeact-mCherry, and nuclear marker (H2B-miRFP670) for later segmentation.

#### Laser-wounding assay

Image acquisition and wounding were performed on a LSM780 confocal microscope (Zeiss), with a 63x, NA 1.4 oil immersion objective (Plan-Apochromat) in a heated stage-top ZILCS incubator (Tokai Hit Co.) at 37°C. Ablation was performed with a Chameleon Vision NLO pulsed laser (Coherent), at 18% power, 10 ms dwell time and 820 nm wavelength. A single circular ROI with 10 pixel diameter (0.38 µm^2^) was ablated at the cortical plane for each cell, and consecutive images acquired with 488 and 564 nm diode lasers (Cobolt).

#### Single-particle fluorescence microscopy

Imaging of single INF2 particles in live cells was performed in TIRF mode using a 100x (NA 1.49, Olympus) oil immersion objective with a pixel binning of 2×2. Bleach-spot triangulation and TIRF 0°-angle were calibrated at the start of every experiment, and TIRF angles were adjusted before each movie. cATP addition and photolysis were performed as for the CaAR assays. Image acquisition was carried out using a 360° rotating TIRF illumination with 488 nm laser intensity set to 100%, acquiring 1 frame every 100 ms. Prior to cATP uncaging, 30 reference frames were acquired.

### Structure prediction and analysis

#### Cryo-EM grid preparation

2.8 µl of sample was applied to glow discharged Quantifoil R2/1 300 Cu mesh holey-carbon grids. Excess sample on the grid was blotted from both sides using a Vitrobot Mark IV (Thermo Fisher Scientific), with a blotting time of 8 s and a blotting force of-20 at 13 °C and 95% humidity. The grids were then plunged in liquid ethane.

#### Cryo-EM data collection and pre-processing

The dataset was collected on a 200 kV Talos Arctica cryo-microscope (Thermo Fisher Scientific) with a Falcon 3 direct electron detector (Thermo Fisher Scientific) operated in linear mode using EPU 1.7. A total of 572 movies were collected with a defocus range of-2.5 to-1.2 μm, a total electron exposure of 57 e^-^/Å^2^ at a pixel size of 1.21 Å in 40 frames.

The dataset was pre-processed on-the-fly using TranSPHIRE version 1.5.13 ^47^. Within TranSPHIRE, beam-induced motion correction and CTF estimation have been performed by UCSF MotionCor2 v1.3.0 ^48^ and CTFFIND4.13 ^49^, respectively. Filament segments were picked using SPHIRE-crYOLO v1.8 ^50^ in filament mode with a boxing distance of 23 pixels (corresponding to 27.5 Å), which resulted in 618,693 filament segments, hereafter referred to as particles.

#### Cryo-EM data processing

A preliminary dataset of INF2-N-bound actin filaments was collected from a sample that was prepared as described above with the only difference that the final buffer contained 5% glycerol and only 0.02% Tween. Filament picking resulted in 444,419 particles which were extracted in a 300 Å^2^ box and imported to CryoSPARC v4.2.1 ^51^. After selection of 2D classes that clearly showed filaments (433,423 particles), a non-uniform refinement was performed using a bare actin filament as an initial volume ^52^. The resulting density clearly showed that INF2-N bound to the actin filament but only at high thresholds. This was used as a starting model for subsequent processing.

For the primary dataset, 618,693 particles were extracted in RELION ^53^ with a box size of 300 Å^2^ and then imported into CryoSPARC. These particles were 2D classified to remove non actin filament particles and the resulting 587,723 particles were helically refined and also non-uniformly refined using the previously obtained INF2-N-decorated actin filament map as an initial reference. Particles that originated from the same filament were grouped together in the same half map for all refinements. These particles were converted back to RELION, where the estimation of beam-induced particle movement was improved by Bayesian Polishing ^54^; and CTF refinements ^55^ were performed to estimate per-particle defoci and to correct for microscope aberrations. After performing a masked 3D auto-refinement in RELION (with solvent flattening Fourier shell correlations (FSCs) enabled), a second round of particle polishing and CTF refinements was done. These corrected particles were imported in CryoSPARC and used for a helical refinement using a helical twist of-166° and a helical rise of 27.5 Å yielding a map of 3.41 Å resolution according to the FSC=0.143 criterion. In this map, the actin-binding site of INF2-N is of high quality but the outer helices are not well resolved. Since the occupancy of INF2-N bound to actin is only around 50-60%, we set out to improve the general density of INF2-N in the central subunit in the box. For this, we created a map comprising only the central INF2-N by fitting an Alphafold-predicted model ^56^ into the density of the helical refined map and then generating a density map from the model at 15 Å using the molmap command in ChimeraX ^57^. A mask was created from this map in CryoSPARC using a dilation radius of 1 and a soft padding width of 12. This mask was then used as a focus mask to subject the particles to a 3D classification in 2 classes, using a class similarity of 0.1. The resulting classes showed one map where the central INF2-N is present and bound to actin (class 0, 334,773 particles) and one class where INF2-N is absent in this position (class 1, 252,950 particles). Particles corresponding to class 0 were then subjected to a helical refinement (final helical parameters: twist-166°, rise: 27.5 Å), and the resulting map (resolution 3.48 Å) showed an improved density for the central INF2-N.

#### Model building

Human protein sequences were obtained from UniProt for inverted formin 2 (INF2, Q27J81), calmodulin 1 (CaM, P0DP23), and β-actin (G8FSY7).

#### INF2-N-actin complex

A published β-actin model bound to phalloidin (PDB: 6T1Y)^58^ was rigid-body fitted in the central region of the filament density using UCSF ChimeraX to generate a starting model for actin. The structure of INF2-N was predicted using AlphaFold2^59^ run on the Raven supercomputer of the Max Planck Computing and Data Facility. After rigid-body fitting the AlphaFold2-predicted INF2-N model into the central subunit of the density, the fitting was further improved by dynamic flexible fitting with Namdinator^60^. The four actin subunits and the central INF2-N were refined by iterative manual model building in Coot^61^ and automated refinement using Phenix real-space refine^62^ with applied geometric restraints. None of the actin subunits were drastically changed from the starting model, only the D-loop and the interacting region with the subunit above had to be adjusted. Due to the high mobility of the outer region of INF2-N, the last 5 helices of INF2-N were rigid body fitted individually and then real-space refined in Phenix. The stretch corresponding to aa 23-31 of INF2-N, which is part of the actin-binding site had to be manually fitted into the density as neither Alphafold2 or Namdinator were able to properly predict its structure.

#### INF2-N interactions with CaM and INF2-DAD

Predictions of complexes between INF2-N (aa 1–269) and CaM (residues 1–149) or INF2-N and INF2-DAD were carried out using a ColabFold v1.5.5 implementation of AlphaFold2 using an MMseqs2-based pipeline. All predictions were run using Google Colab cloud resources under default input parameters.

#### Image analysis and statistics CaAR analysis

Pre-and post-stimulation time-lapse movies from cATP CaAR assays were merged into single BigTiff files using OfflineAnalysis and exported from the software along with a reference image containing the nuclear channel required for segmentation. To handle both the BigTiff datasets and their associated metadata, the Python tifffile library was used. Cell and nucleus outlines were predicted using Cellpose cyto2 and nuclei models, respectively, with diameters empirically set to 150 pixels for cells and 60 pixels for nuclei. To ensure each nucleus was properly assigned to its corresponding cell, the nuclear masks were eroded by 13 pixels (using a disk element), and each nucleus was paired to a cell if its eroded area was fully contained within that cell’s mask boundary.

CaAR measurements made use of three subcellular regions of interest (ROIs): the cytosol, defined as the symmetric difference between the full cell and nucleus masks; the cortex, generated by a 3-pixel dilation of the cell boundary; and the perinuclear region, produced by a 3-pixel dilation of the nucleus boundary. Background intensity levels for each movie were defined as the 0.1st percentile of pixel intensities, which were subsequently subtracted from all pixels. Bleach correction was performed after BG subtraction using a simple ratio approach: for each pixel, the pixel value was multiplied by the ratio of the mean intensity of the first frame divided by the mean intensity of the given frame. Next, the maximum amplitude of CaAR responses and the time point at which this maximum occurred were computed. The relative change in intensity was calculated by dividing all post-stimulation intensities by the average of pre-stimulation values on a per-pixel basis.

#### Single particle fluorescence microscopy (SPFM) analysis

Initial processing of single speckle microscopy datasets was performed using TrackMate (ImageJ plugin), which was configured with the Laplacian of Gaussian (LoG) detector at an expected object radius of 230 nm with subpixel localization. Individual movies required separate threshold adjustments, generally within the range of 180–250. Spot linkage was performed with the Kalman tracker, using a 200 nm initial search radius, a 300 nm maximum search radius, and permitting a maximum gap of two frames. No spots identified by the LoG detector were excluded a priori, and all resulting trajectories were retained for subsequent analysis.

Following TrackMate processing, data were transferred to Python for further analysis. Tracks shorter than three frames were removed, and any missing spots caused by gaps were interpolated by evenly spacing positions between the flanking spots. To avoid overestimating mobility due to detection uncertainties, the mean rate of change in distance from the track’s origin was calculated for each trajectory, rather than using absolute movement. A threshold of 0.03 (empirically determined) served to distinguish between motile and immobile states: if the rate of change between the two preceding points exceeded 0.03, the spot was classified as motile at the current time step. Contiguous motile or immobile segments shorter than or equal to three frames were smoothed to ensure a consistent classification. Since the first two frames required manual adjustment, if movement exceeded 210 nm in the first two frames, these points were designated as motile to accurately capture the state when quick displacement occurred early in a track. This procedure allowed classification of every spot in terms of its mobility status, defining the order, velocity, and duration of motile and immobile segments within each trajectory.

Several practical factors potentially affect these data. Because INF2 activation likely begins already during the 1.5 seconds of photoactivation, a small number of speckles may appear in the first post-stimulation frames; however, significant changes in INF2 behavior generally do not manifest prior to the first post-uncaging image. The TIRF field used for imaging imposes a restricted focal volume, so speckles frequently move in or out of focus, especially if they are actively translocating. This limits the observed lifetimes of tracks, and punctate intensity changes as the z-position varies. A minimum filter for tracks of 400 ms (4 frames) was applied to suppress background fluorescence, and at least three consecutive frames of significant directional displacement were required to classify a speckle as motile. These constraints lead to under detection of very brief elongation events, which may be classified as stationary due to random positional fluctuation matching the characteristic movement threshold. Furthermore, with a 100 ms acquisition rate, processes faster than this timescale may not be captured. Collectively, these factors bias the dataset toward immobile states and against trajectories featuring brief motile phases, but they ensure robust segmentation and analysis of longer-lived speckles and their movement patterns.

#### Colocalization analysis

For colocalization studies, cell ROIs were generated using Cellpose as previously described. After background subtraction, Pearson’s correlation coefficients were computed between the pixel intensities of different fluorescence channels within each ROI to quantify the degree of signal overlap.

#### MRTF quantification

High-throughput quantification of MRTF translocation was performed using immunofluorescence staining in 96-well glass-bottom plates followed by automated imaging on an Olympus ScanR system. Cells were stained for MRTF-A or MRTF-B, F-actin (phalloidin), and nuclei (Hoechst). Automated single-cell analysis was conducted using CellProfiler (v4.2.6) with a Cellpose module (v2.0.5) in an Anaconda environment. Intensities were rescaled from 0-1 (16-bit input). Nuclei were segmented using the Hoechst channel (Cellpose “nuclei” mode), and whole-cell masks were generated using the phalloidin channel (“cyto2” mode). Objects touching image borders were excluded, and nuclei were filtered for size (>150 pixels) and circularity (form factor ≥0.78) to remove dividing or abnormal cells. Only cells with a single associated nucleus were retained. Cytosolic masks were defined by subtracting nuclear from whole-cell masks. MRTF and mNeongreen-INF2 intensities were quantified in nuclear and cytosolic compartments. Image quality was assessed using a focus score metric, and images exceeding a threshold (0.018244) were excluded. Background subtraction was applied using empirically determined values (MRTF-A: 0.003397; MRTF-B: 0.004398). mNeongreen-INF2–positive cells were identified based on cytosolic intensity thresholds derived from INF2 knockout controls (threshold: 0.008305), minimizing false positives (∼1%).

## Statistical analysis and data presentation

Each experiment was repeated at least three times unless stated otherwise. Data presented as boxplots represent measurements aggregated from at least three independent experiments. In each boxplot, the central line indicates the median, the box edges define the interquartile range (IQR; 25th to 75th percentiles), and the whiskers extend to 1.5 times the IQR from the box edges. Notches around the median indicate the 95% confidence interval defined as median±1.57×(IQR/√n).

Time-course data are presented as mean traces of perinuclear or cortical fluorescence intensity (I/I0) plotted against time. Each mean trace was derived from individual cell measurements pooled across at least three independent experiments. The plotted line represents the mean intensity at each time point, and the surrounding shaded area indicates the standard error of the mean (SEM).

## Acknowledgements

S.W and R.J. were members of CiM-IMPRS, the joint graduate school of the Cells-in-Motion Interfaculty Centre, University of Münster and the International Max Planck Research School–Molecular Biomedicine, Münster, Germany. This study was supported by the Deutsche Forschungsgemeinschaft/DFG (**SFB 1557 P15, TRR 422 B01**, **SFB 1348 A01 to R.WS.; SFB 1009 B10 to R.WS and H.S.; SFB 1348 A05 and TRR 422 B04 to M.K.; BI 19998/2-1 to P.B.**), the IZKF Münster (Wed2-022-18 to R.WS; Kr4-003-25 to M.K.), the Royal Society (RSWF\25\R1\1001 to P.B.), the Human Frontier Science Program/HFSP (CDA00070/2017-2 to P.B.), the Max Planck Society (to P.B. and S.R.) and the European Research Council under the European Union’s Horizon 2020 Programme (ERC-2019-SyG, grant no. 856118 to S.R.). This work reflects only the authors’ view and the European Union’s Horizon 2020 research and innovation programme is not responsible for any use that may be made of the information it contains.

## Supplementary figures

**Figure S1.**
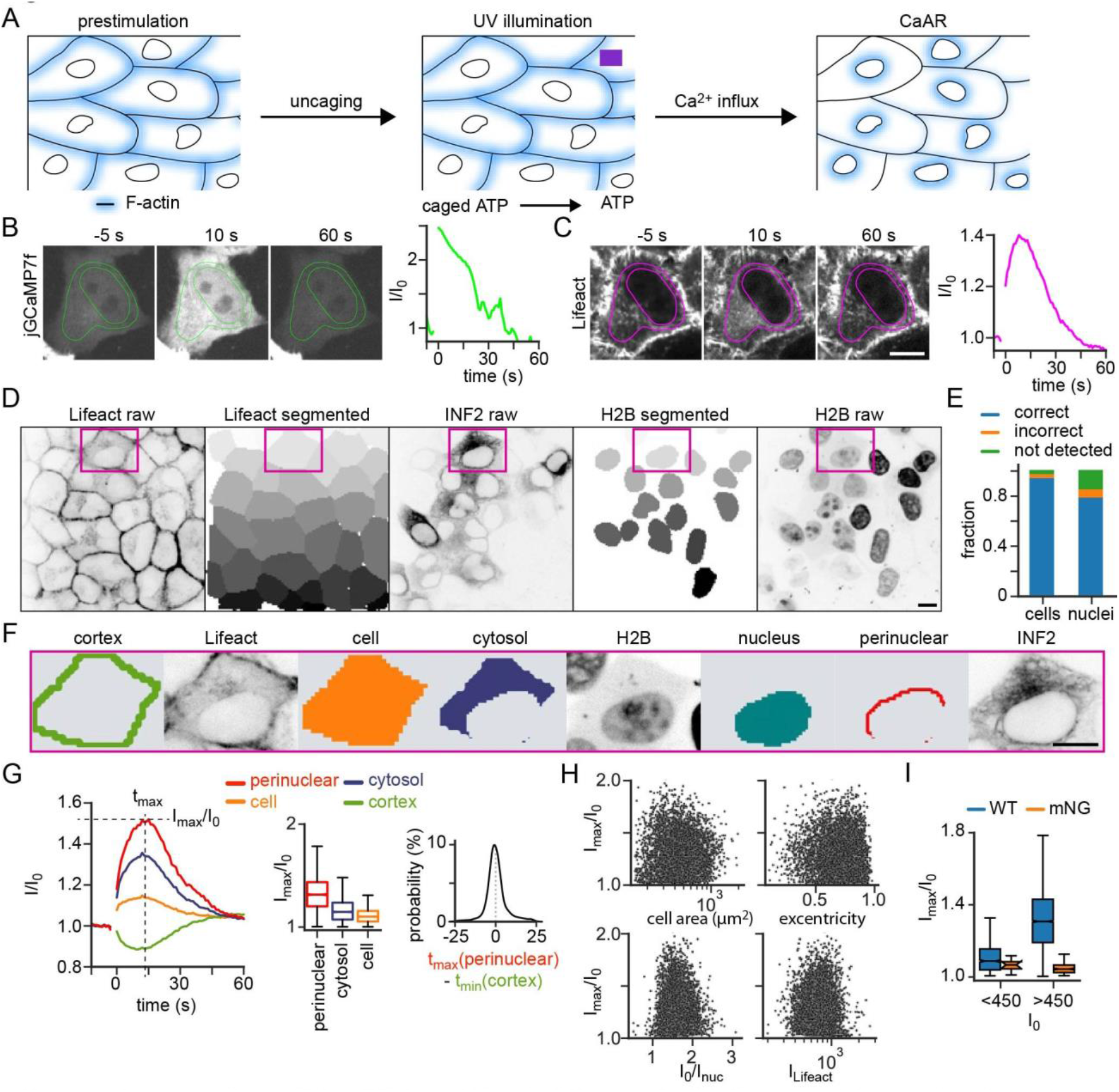
Assay for cellular INF2 activity (CaAR assay). **(A)** Experimental procedure for calcium stimulation via ATP uncaging. **(B, C)** jGCaMP?f (8) and Lifeact-mCh (C) signals in HeLA INF2 KO cells expressing mNG-INF2-CAAX at indicated timepoints before or after uncaging. Exemplary intensity profiles are shown. **(D)** Image analysis pipeline. HeLA INF2 KO cells expressing Lifeact-mCh, mNG-INF2 and miRFP670-H2B were segmented via Cellpose. The Lifeact signal was used to segment cells and the H2B signal to segment nuclei. **(E)** Segmentation fidelity validated by manual assignment. **(F)** Subcellular ROls obtained from the cell in (D). **(G)** Temporal profiles of Lifeact intensities (left, uncaging of ATP at **t** “’ 0) and boxplots of maximal actin reorganization (middle) for ROls indicated in **(F).** Temporal correlation between actin changes at cell cortex and nuclear periphery is shown on the right). **(H)** Scatter plots show lack of correlation between maximal actin reorganization (lmax/I_0)_ and cell area, cell exentricity, perinuclear actin levels (lnuc) and Lifeact-mCh expression (luteact)-**(I)** Boxplots showing minimal mNG-INF2-CAAX expression levels required (>450 grey levels) to obtain actin reorganization. Scale bars: 10 µm.

**Figure S2.**
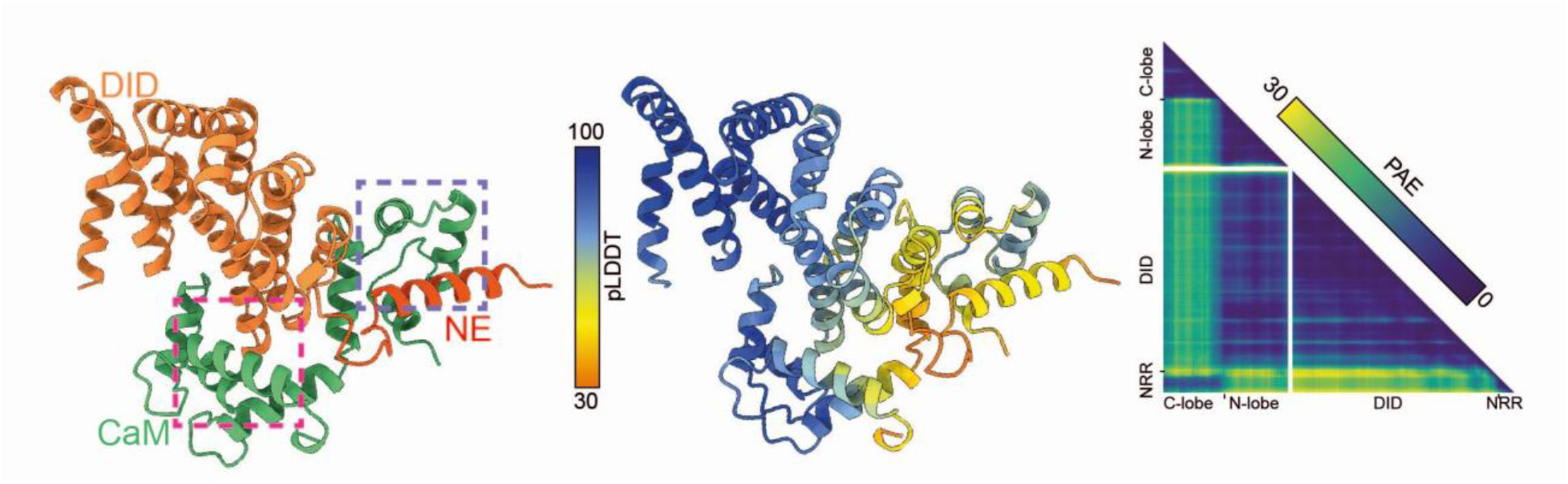
**Interaction of the INF2-N terminus with CaM**. Alphafold 2 prediction with predicted local distance difference test (pLDDT, high confidence for values >70) and predicted alignement error (PAE) of the complex between INF2-N (orange, red) and CaM (green). Boxes indicate interfaces shown in Fig. 2A.

**Figure S3.**
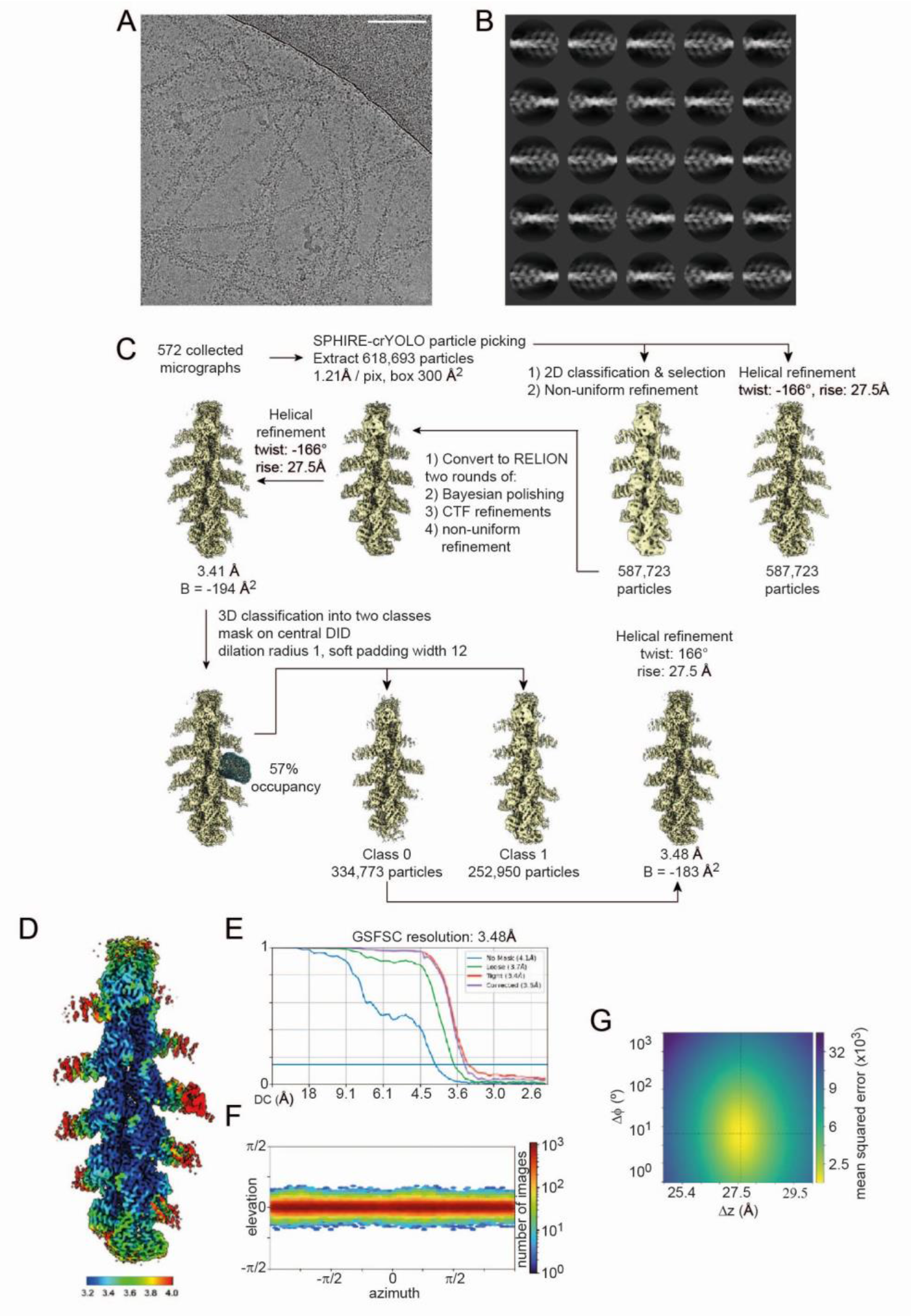
Cryo-EM image-processing of F-actin decorated by INF2-DID. **(A)** Micrograph of INF2-DID-decorated !3-actin filaments frozen in vitreous ice, at a defocus of 2.3 µm. Scale bar: 100 nm. **(B)** Exemplary 2D-class averages obtained with RELION. The box size is 300 A2. **(C)** Image processing pipeline that was used to obtain the different maps. All maps are shown in the same orientation. **(D)** Local resolution estimation of the INF2-DID-decorated F-actin density map calculated in CryoSPARC. The bar shows the range of colors used for each resolution value in A. (E) Fourier-shell correlation calculations between two independent half-maps after applying a loose mask (green), a tight mask (red), no mask (light blue) or a tight mask and correction by noise substitution (purple) of INF2-DID-decorated F-actin particles. The FSC = 0.143 threshold is shown as a light blue line. (F) Angular distribution of the particles used to reconstruct the final cryo-EM map of INF2-DID-decorated F-actin. **(G)** Helical symmetry error surface for the final helical refinement, showing-167° and 27.5 A as the most likely helical twist and rise values.

**Figure S4.**
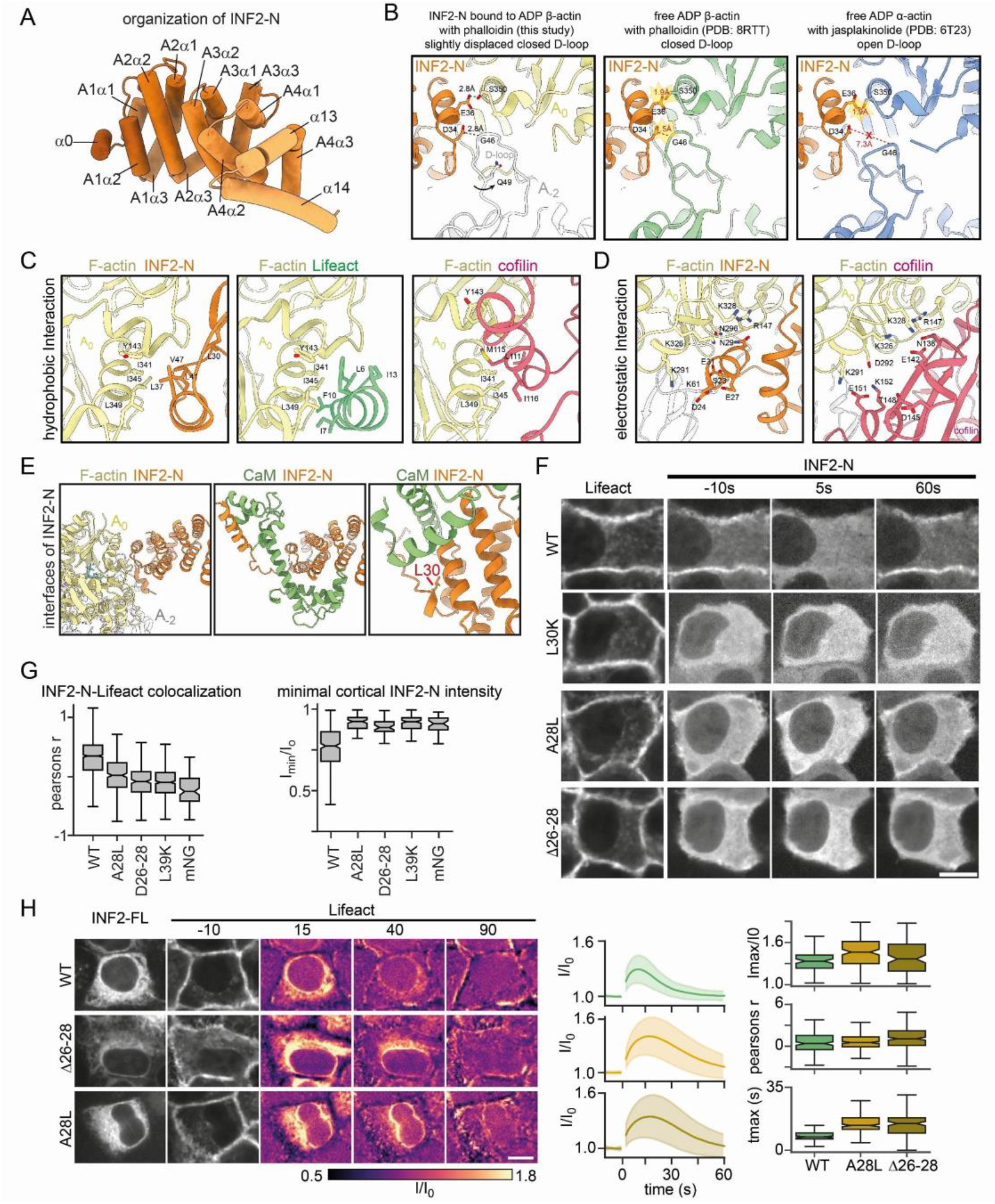
The INF2 N-terminus binds to actin filaments. **(A)** Schematic representation of INF2-N structure. Individual alpha helices (a1-3) for each Armadillo repeat (A1-4) as well as alpha helices in the N-terminal extension (a0) and linking to the DD (a!3/14) are shown. **(B)** Comparison of the D-loop orientation in the cryoEM structure of INF2-N bound to ADP J:l-actin (left) with free phalloidin-bound ADP J:l-actin (middle) and free jasplakinolide-bound ADP a-actin (right). Residue positions favoring the closed D-loop state are indicated. **(C)** Structural comparison of the hydrophobic binding hotspot for the INF2 N-terminus (purple), Lifeact (green, Belyy et al, 2020) and cofilin (red, Huehn et al, 2020). **(D)** Structural comparison of the electrostatic binding hotspot for the INF2 N-terminus (purple) and cofilin (red, Huehn et al, 2020). (E) Comparison of interfaces of INF2-N with CaM and F-actin.The L30 residue is indicated. **(F)** INF2-N-DD cortex association in Hela INF2 KO cells expressing Lifeact-mCh and indicated mNG-INF2-N-DD constructs. (G) Boxplots for the correlation coefficient between Lifeact-mCh and mNG-INF2-N-DD signals at I= 0 and minimal levels of cortical INF2-N-DD association (lmin/lo) after calcium stimulation for the cell lines shown in (F) (n _>_ 70). (H) Actin reorganization upon calcium stimulation in HeLa INF2 KO cells expressing Lifeact-mCherry and indicated mNG-INF2-CAAX constructs. Graphs on the right indicate normalized Lifeact intensity profiles 1/10. Boxplots are shown for maximal levels of actin reorganization (lmax/10, C), the corresponding timepoint (!max, D) and the linear correlation coefficients between Lifeact-mCherry and mNG-INF2 signals at t = 0 (n _>_ 500, N = 3). Scale bars: 5 µm.

**Figure S5.**
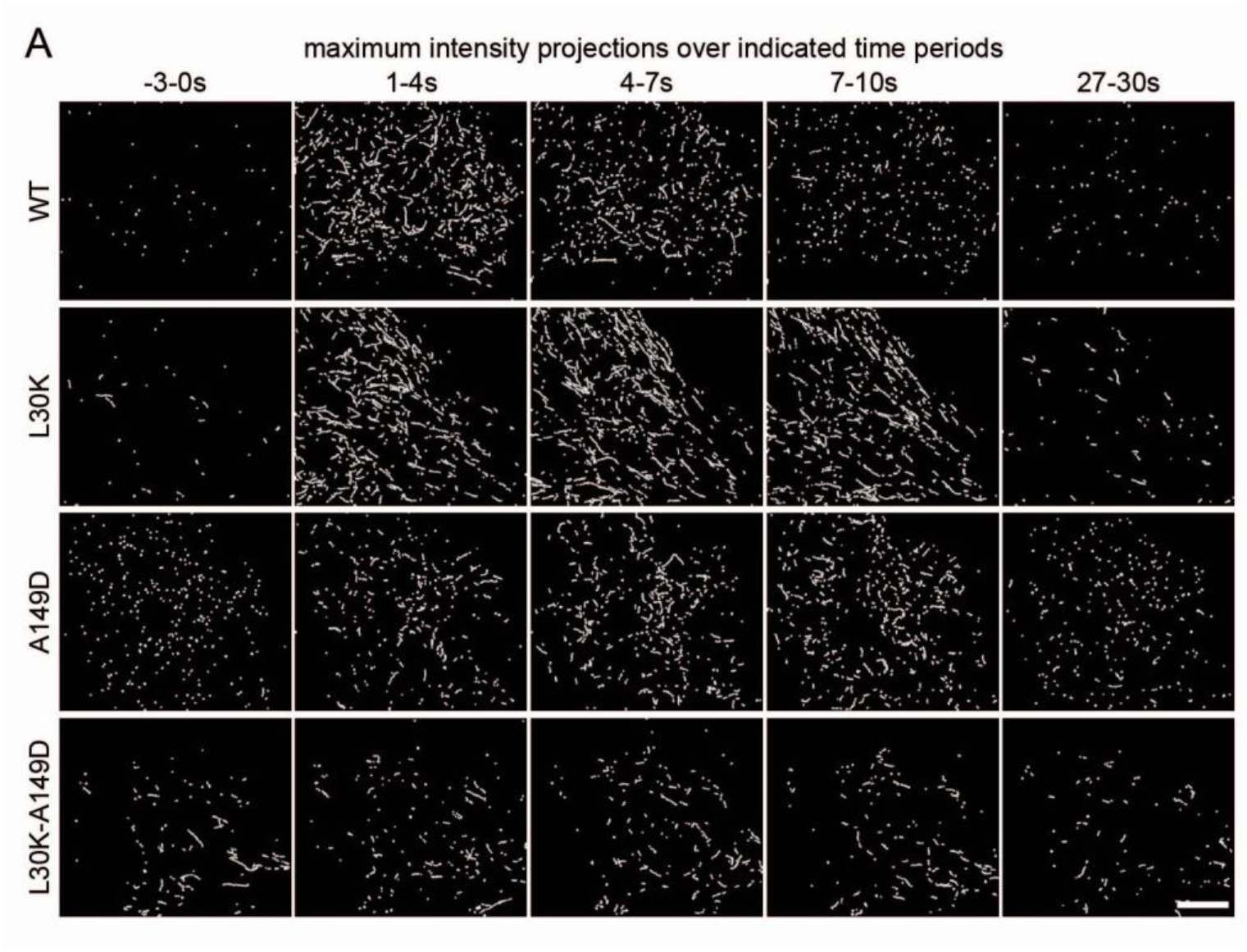
Single particle tracking of INF2 dimers. **(A)** Maximum projection of INF2 speckles identified for indicated periods before and after calcium stimulation. Projections are shown for different regulatory variants depicted in Figure 4. Scale bar: 5 pm.

**Figure S6.**
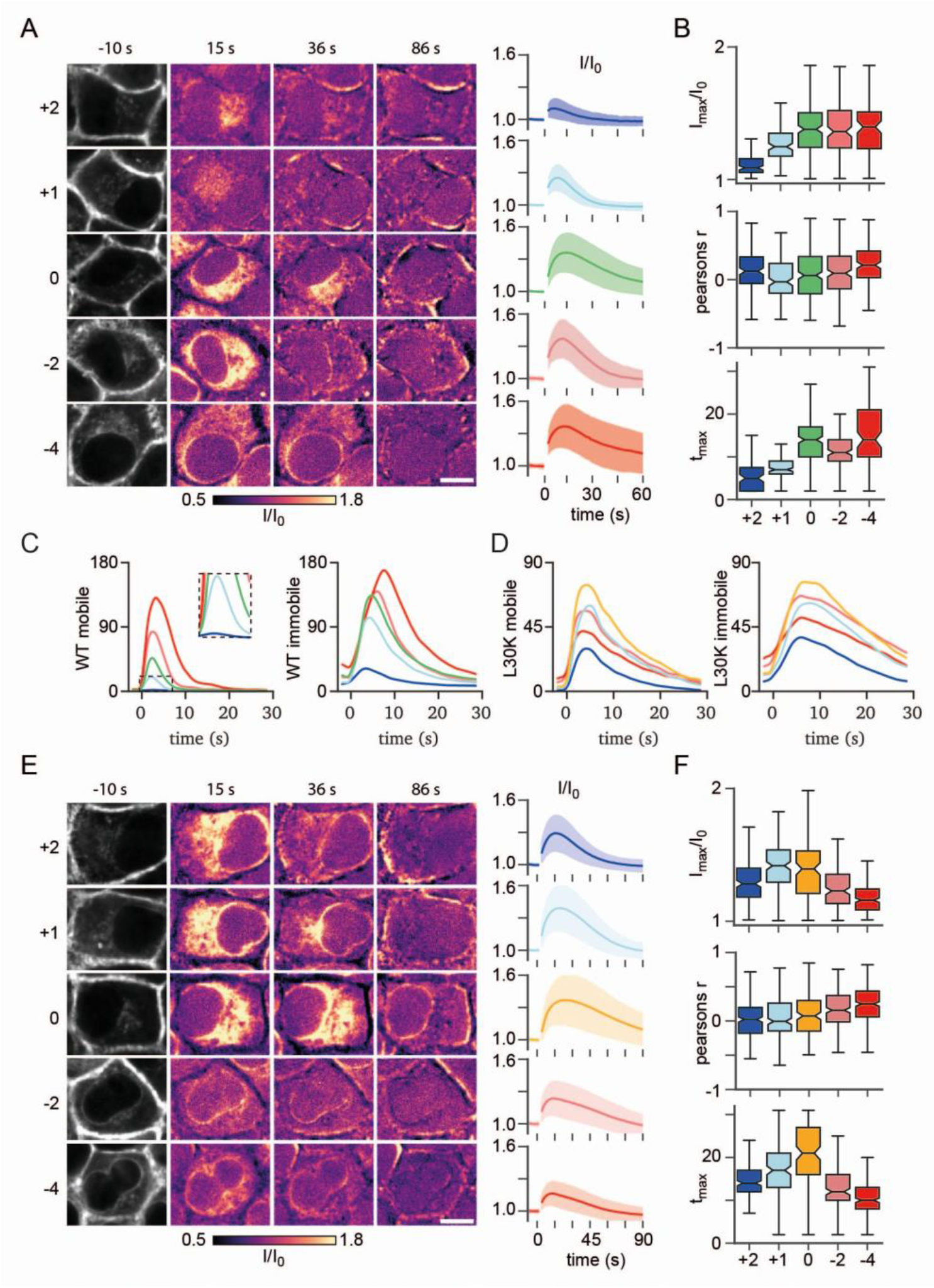
Multi-step re-inhibition of INF2. (A, **B)** Actin reorganization by charge variants of wt mNG-INF2 upon calcium stimulation. Hela INF2 KO cells expressing Lifeact-mCh and indicated mNG-INF2-CAAX constructs were imaged at indicated time points by spinning disk microscopy. Lifeact-mCh images prior to uncaging of caged ATP are shown in grayscale. Images after stimulation show relative change in intensity (fire LUT) compared to the pre-stimulus condition. Graphs on the right indicate normalized Lifeact intensity profiles 1/10 (Mean ± SD, n > 300, N > 2). Scale bar: 5 µm. (C, D) Plots of absolute motile and immobile speckle numbers for indicated charge variants of mNG-INF2 wt (C) and mNG-INF2 L30K (D) (n > 50 cells with > 50k spots, N > 3). (E) Actin reorganization by charge variants of mNG-INF2 L30K upon calcium stimulation. Hela INF2 KO cells expressing Lifeact-mCh and indicated mNG-INF2-CAAX constructs were imaged at indicated time points by spinning disk microscopy. Lifeact-mCh images prior to uncaging of caged ATP are shown in grayscale. Images after stimulation show relative change in intensity fire LUT) compared to the pre-stimulus condition. Graphs on the right indicate normalized Lifeact intensity profiles 1/10 (Mean ± SD, n > 300, **N** > **2).** Scale bar: 5 µm. (F) Boxplots for maximal levels of actin reorganization (lmax/10), the corresponding time point (tmaxl and linear correlation coefficients between Lifeact-mCh and mNG-INF2 signals at **t** = O for the cell lines shown in (E).

**Figure S7.**
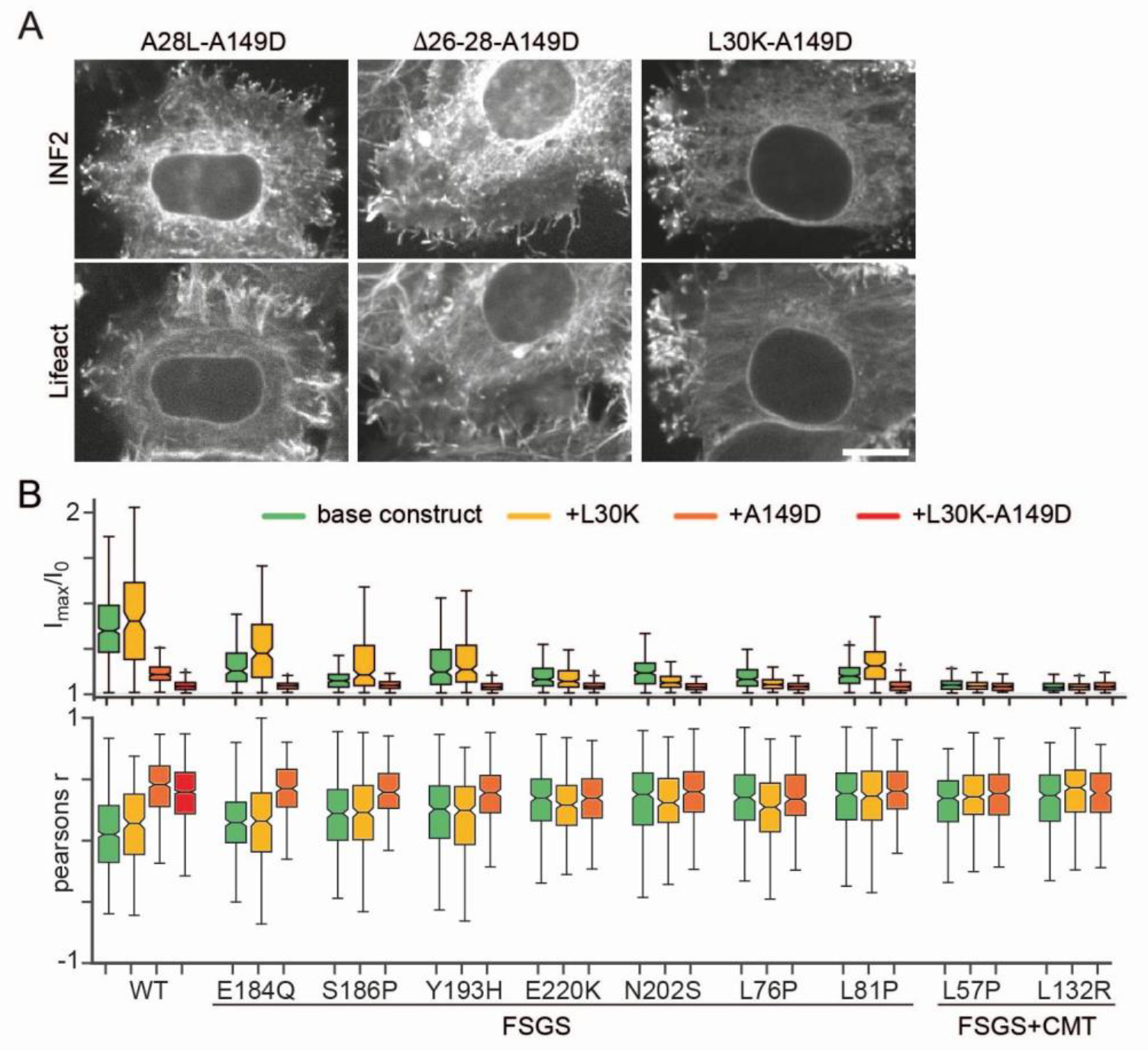
Physiological consequences of INF2 deregulation and model of INF2 activity cycle. **(A)** Formation of filopodia in HeLa INF2 KO cells espressing indicated fully activated mNG-INF2 variants. mNG-INF2 localizes to the tips of actin filled fillopodia. Scale bar: 5 pm. (B) Boxplots for maximal levels of actin reorganization after calcium stimulation (l_m_ax/lo) and correlation coefficients (pearsons r) between Lifeact-mCh and mNG-INF2 signals prior to stimulation. Values are shown for HeLa INF2 KO cells expressing indicated mNG-INF2-CAAX variants associated with FSGS or FSGS+CMT in patients, or the respective disease variants combined with additional regulatory mutations (n > 50, N = 3).

## Supplementary movie legends

**Movie S1**. Actin reorganization by INF2 upon calcium stimulation. HeLa INF2 KO cells expressing Lifeact-mCherry and indicated mNG-INF2-CAAX constructs (or mNeonGreen - mNG as control) were imaged by spinning disk microscopy. Movie shows Lifeact-mCherry signal in grayscale. Time in seconds. Corresponds to Fig. 1B.

**Movie S2.** Animation illustrating the small change in the actin D-loop structure upon binding of INF2-N. Corresponds to Fig. S4B.

**Movie S3**. Animation illustrating the hydrophobic and electrostatic binding interfaces of INF2-N on F-actin. Corresponds to Fig. 3D.

**Movie S4.** Single particle imaging of indicated mNG-INF2-nCAAX variants by TIRF microscopy. Calcium stimulation via uncaging of cATP occurs at t = 0 s. Time in seconds, Scale bar 5 µm. Corresponds to Fig. 5A.

**Movie S5.** Formation of filopodia in HeLa INF2 KO cells expressing Lifeact-mCherry and indicated mNG-INF2 variants at different times after PM wounding (wound positions indicated with asterisks, wounding at t = 0 s). Time in seconds. Scale bar 5 µm. Corresponds to Fig. 7B.

